# Foliar water uptake as a source of hydrogen and oxygen in plant biomass

**DOI:** 10.1101/2020.08.20.260372

**Authors:** Akira Kagawa

## Abstract

Introductory biology lessons around the world typically teach that plants absorb water through their roots, but, unfortunately, absorption of water through leaves and subsequent transport and use of this water for biomass formation remains a field limited mostly to specialists. Recent studies have identified foliar water uptake as a significant but still unquantified net water source for terrestrial plants. The growing interest in the development of a new model that includes foliar uptake of liquid water to explain hydrogen and oxygen isotope ratios in leaf water and tree rings requires a method for distinguishing between these two water sources. I therefore devised a method utilizing two different heavy waters (HDO and H_2_^18^O) to simultaneously label both foliar-uptake water and root-uptake water and quantify their relative contributions to plant biomass. Using this new method, I here present evidence that, in the case of well-watered *Cryptomeria japonica*, hydrogen and oxygen incorporated into new leaf cellulose in the rainy season derives mostly from foliar-uptake water, while that of new root cellulose derives mostly from root-uptake water, and new branch xylem is somewhere in between. Abandoning the assumption that these elements are supplied from soil water alone may have vast implications in fields ranging from isotope dendroclimatology, silviculture, to biogeochemistry.

---

The concept that terrestrial plants absorb water through their roots and release water into the air through their leaves is common knowledge (McElrone et al., 2013) that has formed the basis for models for irrigation, atmospheric sciences, and isotope dendroclimatology, but, in recent decades, the underlying assumptions of these models have started to fall apart in response to growing evidence that plants can absorb much of their water through their leaves (Eller et al., 2013; Goldsmith et al., 2013; Dawson & Goldsmith, 2018; Berry et al., 2019; Schreel & Steppe, 2020). Water in both liquid and vapour phases can be absorbed through foliar trichomes, cuticles, hydathodes, and stomata (Fernandez et al. 2014; Ritpitakphong et al., 2016; Fernandez et al., 2017; Schreel et al. 2020). Especially, physicochemical properties of trichomes, epidermis and other leaf organs play an important role for absorption of water into leaf (Fernandez et al. 2014; Azuma et al.2020; Schreel et al. 2020).

The first indications that the general model was too simplistic came in the 1980s from studies of fractionation of water in plant leaves (Leaney et al., 1985; Yakir et al., 1990). The ratio of heavy isotopes (δD and δ^18^O) in leaf water tends to be elevated during transpiration because water molecules composed of heavy isotopes (HDO and H_2_^18^O) evaporate slower from leaf surfaces than H_2_^16^O. Although the influence of this enrichment on a number of biogeochemical processes and the resulting ratios of oxygen isotopes in the atmosphere—including isotopes in carbon dioxide, a potential indicator of terrestrial gross primary productivity (Farquhar et al., 1993; Farquhar & Lloyd, 1993; Cuntz et al., 2003; Welp et al., 2011) and isotopes in molecular oxygen, which contributes to the Dole effect (Dole et al., 1954; Luz et al., 1999; Hoffmann et al., 2004)—is now well accepted, the fate of enriched leaf water within plants is less understood. Modifications of the “Craig-Gordon” model (Craig & Gordon, 1965; Dongmann et al., 1974) have tended to overestimate leaf water enrichment, and have been further developed into two mechanistic models: the discrete-pool model and the Péclet model. The discrete-pool model views leaf water as two distinct fractions: evaporatively enriched water in the lamina and unenriched source water in the veins (Leany et al., 1985; Yakir et al., 1989; Yakir et al., 1990; Yakir et al., 1994). The Péclet model takes into account both advection of vein water due to transpiration and backward diffusion of enriched water from evaporative sites, thus generating an isotopic gradient from veins to the evaporation site (Farquhar & Lloyd, 1993; Farquhar & Gan, 2003). However, despite huge efforts to further revise these models, they are still insufficient to describe observed isotopic distributions, and researchers have begun to look for creative solutions (Cernusak et al., 2016). For example, Helliker & Richter (2008) reported a discrepancy between observed oxygen isotope enrichment of tree-ring cellulose above source water (*∆*^18^O_c_) and *∆*^18^O_c_ values predicted from the Péclet model (Barbour & Farquhar, 2000), and attributed the discrepancy to convergence of tree-leaf temperatures in subtropical and boreal biomes. However, there is a latitudinal gradient of precipitation with lesser precipitation and humidity at higher latitude (Baumgartrer & Reichel, 1975). Perhaps some discrepancies between current models and observations can be explained by foliar uptake of liquid water (the net uptake of liquid water through the surface of wet leaves) in addition to root water uptake.

In the past, water in roots and stems was commonly assumed to have the same isotopic composition as soil water (Gonfiatini et al., 1965; Wershaw et al., 1966; Dawson & Ehleringer, 1991). However, in light of current research showing that foliar water uptake provides plants with a significant secondary water source (Eller et al., 2013; Goldsmith et al., 2013; Dawson & Goldsmith, 2018; Berry et al., 2019; Schreel & Steppe, 2020), we should expect isotopic differences between precipitation and soil water (Treydte et al., 2014) to be reflected in different parts of plants (Zhao et al., 2016; Hannes et al., 2020). While foliar vapour exchange with unsaturated air cannot cause reverse advectional flow of water, foliar-uptake liquid water can. Since foliar water uptake—which includes both foliar vapor exchange and foliar-uptake of liquid water—is associated with reversals in sap flow, we might hypothesize the existence of a gradient in the ratio of foliar-uptake water to root-uptake water (FUW/RUW), with the ratio decreasing across tissue water from leaves to roots, and FUW even reaching the roots in some cases (Eller et al., 2013; Cassana et al., 2016). If this gradient is then reflected in biomass, the results of Helliker & Richter’s (2008) study could be explained more simply by the correlation of latitude with precipitation and humidity (Baumgartner & Reichel, 1975). Adequate assessment of this possibility requires investigating the contribution of FUW and RUW to the isotopic gradient along the stem as well as in the roots and leaves, and in both tissue water and biomass.

Understanding how leaf water and cambial water isotope signals are then incorporated into tree-ring cellulose is important for applications such as reconstruction of paleoclimate and analysis involving plant responses to environmental changes (Gessler et al., 2014). In East Asia, archaeological research has come to rely more and more on tree-ring oxygen isotopes for isotopic cross-dating (Roden, 2008; Nakai et al., 2018; Loader et al., 2019; Seo et al., 2019; Choi et al., 2020; Nakatsuka et al., 2020) and provenancing of wood (Kagawa & Leavitt, 2010). The precision of these methods depends on within-site coherence and mechanistic understanding of tree-ring δ^18^O. Isotopic mass balance model of leaf water predicted that significant amount of atmospheric vapour can enter the leaf (Farquhar & Cernusak, 2005), and incorporation of the vapour into plant biomass through stomata is assumed by the model (Roden et al. 2000). However, relative contributions of foliar-uptake vapour and root-uptake water to plant biomass remained unknown. The idea that foliar-uptake liquid water, in addition to atmospheric vapour, may be incorporated into plant biomass has only received serious attention in recent years. For example, Lehman et al. (2018; 2019) used water mist depleted in ^18^O to demonstrate how FUW can be assimilated into sugars formed during photosynthesis in wet and humid climates, but left open the question of FUW assimilation into tree rings. It is therefore vital that we investigate similar possibilities for tree-ring cellulose in climates with a wet growing season.

According to Köppen’s climate classification, the Japanese mainland is characterized by a “humid subtropical climate” (Kottek et al., 2006), in which the summer monsoon brings a marked rainy season (commonly called the “plum rain”, 梅雨, in countries in this region). During the rainy season, overcast conditions with photosynthetic photon flux density (PPFD) of less than 1000 μmol m^−2^ s^−1^ are much more frequent than sunny conditions with PPFD around 2000 μmol m^−2^ s^−1^ (Miyashita et al., 2012). Rice leaves, for example, stay wet for about 70% of this period (Yoshida et al., 2015; Hiromitsu Kanno, unpublished data), and reported durations of leaf wetness in forests are only slightly less (Klemm et al., 2002, Binks et al. 2020). In Tsukuba, where this study was conducted, an average rainy season lasts from June 8 to July 21 (Japan Meteorological Agency, https://www.jma.go.jp/), which corresponds to the latter half of the earlywood formation period and includes the summer solstice when radial tree growth is at a maximum (Kagawa et al., 2005; Rossi et al., 2006).

With the requirement for leaf wetness during the growing season met, we might suspect that foliar water uptake will have an especially strong impact on hydrogen and oxygen isotope ratios in trees in Japan and surrounding regions. This suspicion grows if we consider that, according to statistical isotope dendroclimatology data from various parts of the world, tree-ring δ^18^O series from regions with relatively wet growing seasons tend to show both higher correlation to early-summer humidity and precipitation, and higher within-site coherence. For example, oxygen isotope ratios of tree rings from Japan and Korea—including tree-ring δ^18^O series for Japanese cedar (*Cryptomeria japonica*) with some of the highest known degrees of coherences in the world—show very high correlations to precipitation and humidity during the rainy season (Nakatsuka et al., 2004; Li et al., 2015; Seo et al., 2019; Nakatsuka et al., 2020). In the Pacific Northwest, which has a similarly wet growing season, foliar water uptake has been directly observed in coastal redwoods (*Sequoia sempervirens*) with up to 6% of leaf water originating from the previous night’s fog (Burgess & Dawson, 2004), and tree rings of coastal redwood also show strong coherence at both inter-annual and intra-annual levels (Roden et al., 2008; Roden et al., 2009; Roden et al., 2011). In contrast, in regions with a Mediterranean climate, for example, precipitation and humidity reach a minimum in the middle of the growing season (Kottek et al., 2006), and tree-ring δ^18^O shows weaker coherence (Konter et al., 2014; Shestakova et al. 2014).

This is understandable if we consider that isotope ratios of rainwater (and atmospheric vapour) are strongly coherent from catchment to regional scales (Miyake et al., 1968; Ingraham, 1998; Treydte et al., 2014; Baker et al., 2015), whereas isotope ratios of soil water are less coherent because of the heterogeneous nature of soil (Dawson, 1993; Tsujimura & Tanaka, 1998; Treydte et al., 2014). Although foliar water uptake is an important water source for Mediterranean trees during dry summer (Fernandez et al. 2014), relative contribution of FUW to hydrogen and oxygen of plant biomass should be smaller in such areas compared to regions with wet and humid growing seasons. We can therefore hypothesize that tree-ring hydrogen and oxygen isotopic series with higher contributions of FUW may show stronger coherence across a study site.

Our current mechanistic understanding of the assimilation of FUW into tree rings has been limited by lack of effective tools to study this phenomenon (Berry et al., 2019). Authors including myself have suggested that use of artificially enriched tracers in the forms of ^13^CO_2_, HDO, and H_2_^18^O instead of natural isotope tracers may provide a clearer picture of the physiological processes between photosynthetic incorporation of isotopes and their deposition into tree rings (Roden et al. 1999a; Helle & Panferov, 2004; Kagawa et al., 2005; Kagawa et al., 2006a). I have therefore devised a novel dual-isotope labelling method utilizing HDO and H_2_^18^O (Fig. 1), which are particularly appealing because hydrogen and oxygen isotopes behave similarly in the environment (Craig & Gordon, 1965) and treering isotope ratios of hydrogen and oxygen are explained by the same mechanistic model with different fractionation factors (Roden et al., 2000; Gessler et al., 2014). The dual-isotope labelling method allows separate labelling of foliar- and root-absorbed water with heavy hydrogen water and heavy oxygen water.

**Fig. 1.**
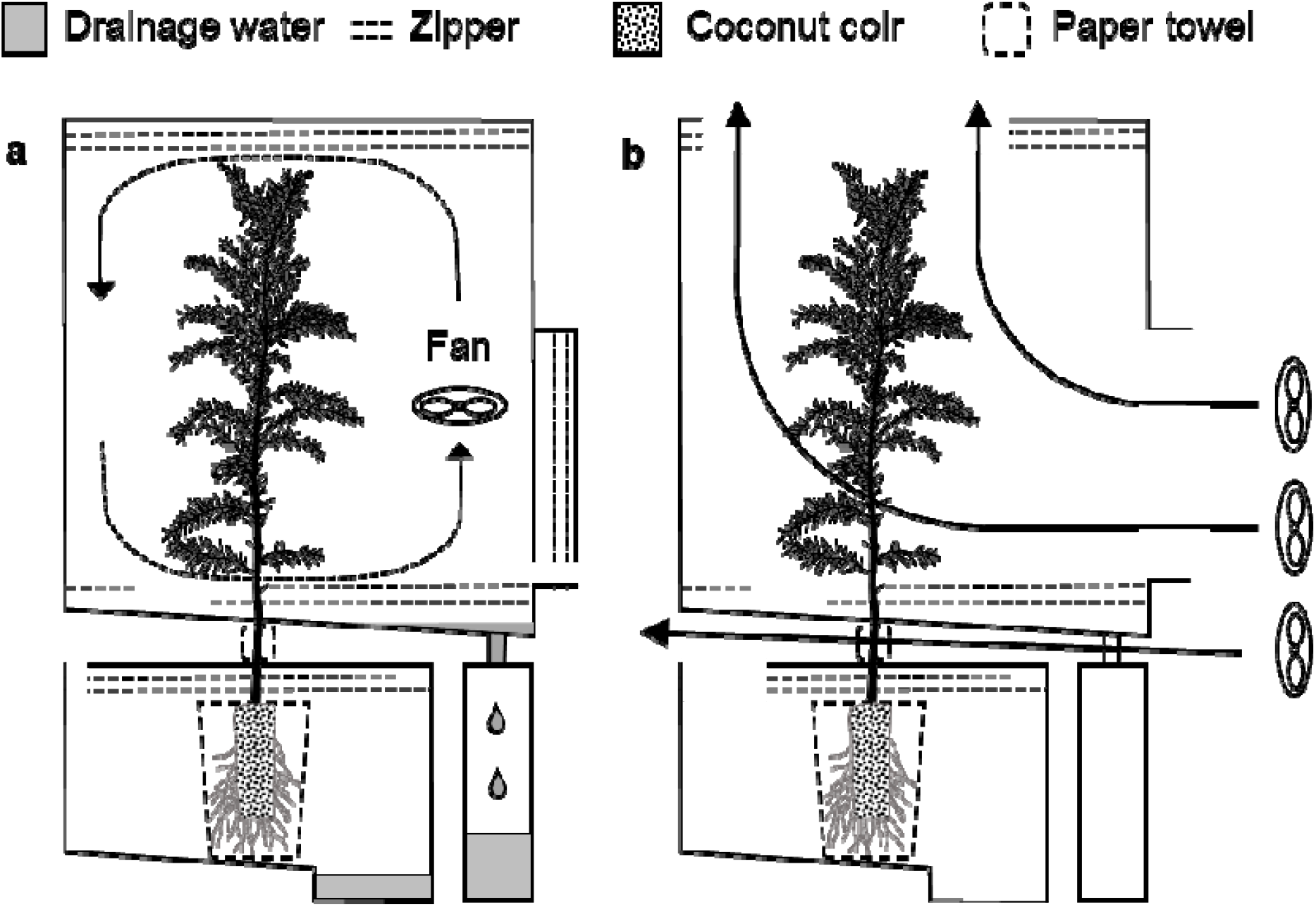
Chambers for dual-isotope pulse-labelling. (a) To label O/H trees (Trees A, B, and C), needles were sprayed with heavy oxygen water (H_2_^18^O) and soil was watered with heavy hydrogen water (HDO). (b) To label H/O trees (Trees D, E, and F), needles were sprayed with heavy hydrogen water and soil was watered with heavy oxygen water. After labelling, chambers were exposed to gentle wind so that vapours potentially leaking from chambers would not enter other chambers.

There are three immediate questions I hope this dual labelling method can help to answer: (1) What is the rate of foliar water uptake and when does it reach equilibrium? (2) Does the FUW/RUW ratio differ across needles, phloem+epidermis, xylem and roots? (3) What percentage of hydrogen and oxygen in cellulose and dry organic matter originates from FUW? In order to answer these questions, I chose Japanese cedar as my subject. Japanese cedar is one of the most frequently studied trees in (isotope) dendroclimatology research in Japan, while their North American siblings in the sequoia genus (family Cupressaceae) are similarly well-known in research. For example, redwood is known as the tallest tree species on earth, and even with ample soil moisture, increasing leaf water stress due to gravity and path length resistance is likely to limit its height (Koch et al., 2004), unless foliar water uptake can provide a second source of water (Burgess & Dawson, 2004). Likewise, foliar water uptake may play an important role for Japanese cedar, which achieves the tallest height among Japanese tree species.

From 16 July at 18:00 through 19 July at 18:00, during the rainy season of east-central Japan, I pulse-labelled one “O/H” group of three saplings by spraying needles with heavy oxygen water and watering roots with heavy hydrogen water, and a second “H/O” group of three saplings by spraying needles with heavy hydrogen water and watering roots with heavy oxygen water (Fig. 1). Two control saplings were supplied only tap water. By analysing the subsequent hydrogen and oxygen isotopic signals of the drainage water and of the tissue water and biomass in each organ, I herein attempt to shed light on the absorption, transport, and assimilation of foliar-uptake water into the structural components of trees.

## 2 Results

### 2.1 Preliminaries

In Tsukuba, the rainy season of 2019 lasted from June 7 to July 24 (Japan Meteorological Agency, https://www.jma.go.jp/), which puts our pulse-labelling period (July 16 to 19) near the end of the rainy season. The weather during the pulse-labelling period (dotted square in Extended Data Fig. 1) was cloudy, and I was satisfied that foliar water uptake was likely. The four days following the pulse-labelling had low solar radiation with occasional rain and July 24 onward saw a series of sunny days with temperature exceeding 30 ◻. Transpiration was proportional to PPFD, and, trees transpired an average of 53 g each on sunny days and 3-35 g each on rainy/cloudy days (Extended Data Table 1).

Preliminary analysis of heavy water gave the following results, which were used in subsequent sample analysis: Heavy hydrogen water had ^18^O concentration within natural range and deuterium concentration of 1.1%, the equivalent of a deuterium concentration 72 times higher than natural water. Heavy oxygen water had deuterium concentration within natural range and ^18^O concentration of 2.3%, 12 times higher than natural water (Extended Data Table 2). Based on these values, a 960‰ increase in excess δD translates to a 1% increase in τH, and a 140‰ increase in excess δ^18^O translates a 1% increase in *τ*_O_.

Analysis of the control trees showed no significant increase over reference of δ^18^O in needle and δ^18^O and δD in root water (t test, one tailed, *P* > 0.05). There was a significant increase in δD of needle water (24.4‰, *P* = 3.1×10^−5^), but still within natural variation (±26‰) according to Cernusak et al. (2016). I therefore concluded that any cross-chamber contamination was within acceptable limits.

### 2.2 Isotopic signals of FUW and RUW in tissue water

Heavy water concentrations in needles and roots showed similar trends in both an O/H sapling (Fig. 2a) and an H/O sapling (Fig. 2b). Needles sampled during the labelling period showed the highest water content at the upper part (62.2 ± 1.2 %, *n* = 10), followed by middle (57.0 ± 5.8 %, *n* = 10) and lower parts (51.2 ± 3.0 %, *n* = 10), as previously observed in mature trees of this species (Azuma et al., 2016; Azuma et al. 2020), however, the concentration of FUW varied little among upper, middle, and lower needles. Almost half of needle water was replaced by foliar-uptake heavy water (heavy FUW) within 24 hours of heavy water exposure. Immediately after the last heavy water spray (18 July, 18:00), heavy FUW in needles started decreasing, and within 48 hours of opening each chamber on 19 July, 18:00 (Fig. 1b), the heavy FUW signal in needles was completely lost (21 July, 18:00).

**Fig. 2.**
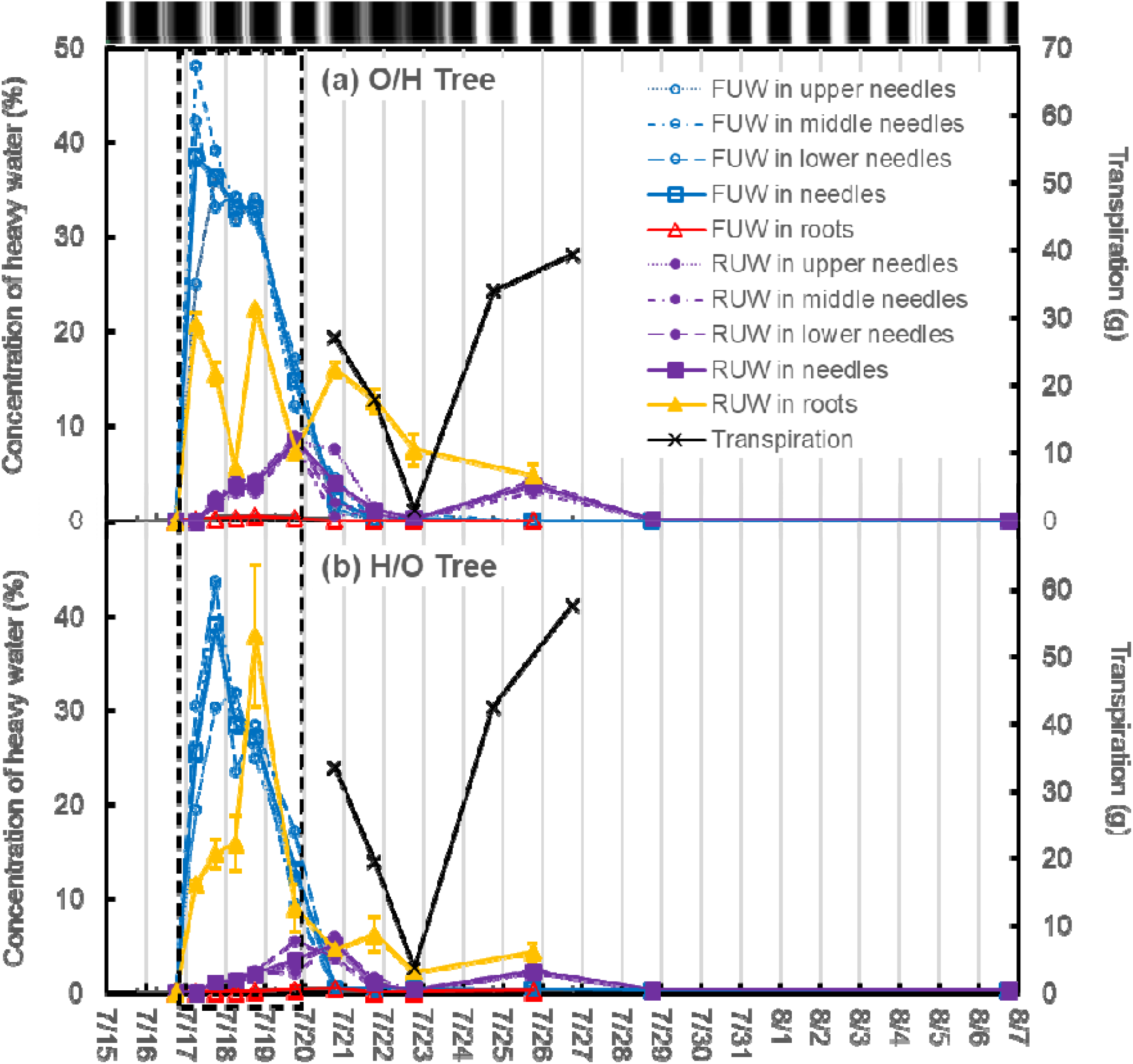
Absorption and transport of heavy foliar-uptake water (FUW) and heavy root-uptake water (RUW) within trees. (a) Concentrations of heavy water in needles and roots of an O/H tree. Needles and roots of this tree were exposed to heavy oxygen water and heavy hydrogen water, respectively. (b) Concentrations of heavy water in needles and roots of an H/O tree. Needles and roots of this tree were exposed to heavy hydrogen water and heavy oxygen water, respectively. Labelling with heavy water was conducted from 16 July at 18:00 to 19 July at 18:00 (dashed box). Heavy FUW signals in needles spiked dramatically during labelling, but disappeared within four days of the last labelled spray, while heavy RUW signals were more erratic and somewhat less pronounced, but still clearly present in both roots and needles seven days after labelling. For reference, the diurnal patterns of PPFD in Extended Data Fig. 1 are illustrated by the grey colour map at the top of the figure. Error bars represent SD of replicate measurements, except for FUW in roots of (a) O/H tree sampled at 06:00 and 18:00 on 18 July, and at 18:00 on 19 July, for which only one data per sample was available.

Root-uptake heavy water (heavy RUW) was first detected in needles 24 hours after the first heavy water administration (17 July, 18:00) and reached a maximum (19 July, 18:00 – 20 July, 18:00) at around the end of the labelling period. Heavy RUW in needles continued to decrease from 20 July, 18:00 until 22 July, 18:00, which corresponds to a rainy period with low PPFD. Then, from 24 July, trees responded to a series of sunny days with increased transpiration. Quite unexpectedly, heavy RUW reappeared strongly in needles on 25 July 18:00, but by 28 July, 18:00, the signal was completely lost.

Heavy RUW in roots showed larger short-term fluctuations than heavy RUW in needles, but roots sampled from different locations showed similar values, suggesting large temporal change and small spatial change in heavy RUW concentrations. Concentration of heavy RUW in roots reached maximum at the end of heavy water administration (18 July, 18:00) then gradually decreased.

Very small but significant amounts of heavy FUW were detected in all root samples of the O/H and H/O trees, except for O/H roots of 20 July, 18:00 (t-test, one tailed, *P* < 0.032). The highest excess δ^18^O and δD values observed in O/H and H/O roots were 51.5‰ and 232‰, respectively.

Newly formed biomass, such as needles, wood and roots, carries time-integrated FUW/RUW signature of local water at the cambium (Roden et al. 2000; Barnard et al. 2007; Gessler et al. 2007). Overall, the time-integrated peak area ratio of heavy FUW to RUW (*∆*^18^O_w_/∆D_w_ for O/H tree and ∆D_w_/*∆*^18^O_w_ for H/O tree in Fig. 2) in tissue water across different organs, ranged from 4.8 in the needles to 0.02 in the roots when averaged over O/H and H/O trees (forming a gradient qualitatively expressed in the left side of Fig. 3).

**Fig. 3.**
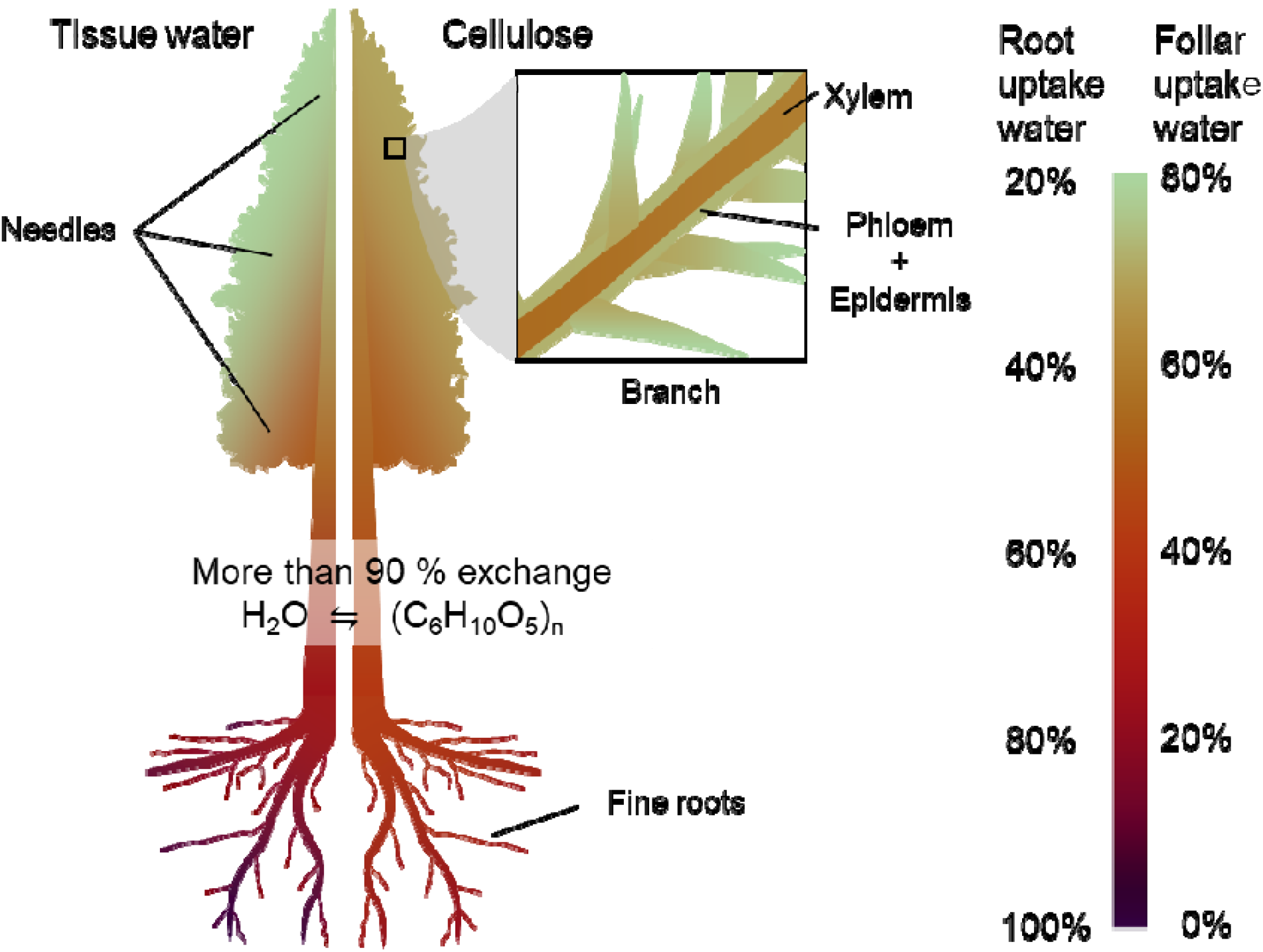
Origins of hydrogen and oxygen in tree biomass formed under rain conditions. During the first three days of the experiment, which were rainy, leaf water was mostly composed of FUW, and root water was mostly composed of RUW. Stem water was estimated to be around half FUW and half RUW (left side of the tree figure). FUW and RUW concentrations in local tissue were reflected in the hydrogen and oxygen of local biomass. About 90% of hydrogen and oxygen in sugars should have exchanged with local water at the cambium. Thus, most of the hydrogen and oxygen incorporated into new leaf cellulose was derived from FUW, while that of new root cellulose was derived mostly from RUW, and that of new branch xylem was derived from roughly equal parts of FUW and RUW.

### 2.3 Isotope ratios of drainage water

Less than half of the drainage water from the aboveground chambers originated from the heavy water sprayed to needles (Extended Data Table 3). About half of the drainage water was natural water originating from inside the trees and soil and a small fraction (4-8%) was heavy water administered to the ground chamber.

About one third of drainage water from the ground chambers consisted of heavy water administered to the soil, and the rest was natural water. The absence of heavy FUW (Excess δ^18^O of O/H tree ground drainage water was 9.1‰, and excess δD of H/O tree ground drainage water was 2.3‰) further indicates that no cross-chamber contamination occurred (Fig. 1).

### 2.4 Heavy FUW and RUW ratios in cellulose

Trends in the ratios of FUW to RUW assimilation into cellulose were examined through analysis of heavy water percentage (τ) in α-cellulose (Extended Data Table 4), where assimilation of both HDO and H_2_^18^O into cellulose was taken into account. ANOVA tests showed no significant difference between O/H and H/O treatments on heavy water percentage (τ) in α-cellulose extracted from needles, phloem+epidermis, xylem, and roots (*P* = 0.40), nor a significant difference in τ between (1) needles and (2) phloem+epidermis (*P* = 0.63). However, (1) needles, phloem, and epidermis, (2) xylem, and (3) roots were significantly different from each other (*P* = 1.8×10^−10^). I was worried that, compared to samples from July 2019, samples from January 2020 may contain lesser label because biomass formed during the rest of the growing season dilutes the labelling signal. However, dilution of the label by the non-labelled biomass was not so large, because growth slows down after the end of the rainy season, and hydrogen and oxygen isotope ratios of cellulose extracted in January showed much higher isotope ratios than natural abundance, except for the roots sampled in January, which was formed from stored carbohydrates (Extended Data Table 6).

Average *τ*_Cell_ of FUW decreased in the order of (1) needles, phloem, and epidermis, (2) xylem, and (3) roots, decreasing with distance from needles (Extended Data Table 4). Similarly, average *τ*Cell of RUW decreased in the order of (1) roots, (2) xylem, and (3) needles, phloem, and epidermis, decreasing with distance from roots. The ratio of assimilated FUW/RUW in α-cellulose ranged from a ratio of 2.2 in the needles to 0.2 in the roots (forming a gradient qualitatively expressed in the right side of Fig. 3).

## 3 Discussion

### 3.1 Foliar and root water uptake

The first question I aim to address in this paper concerns the rates of foliar water uptake and root water uptake and the process of absorption. In this study, even though water content increased with needle elevation within saplings—which we would expect since needles near the tops of Japanese cedars show the most succulence and have the most transfusion tissue for storing water (Azuma et al., 2016, Azuma et al. 2020)—the concentration of FUW varied little among upper, middle, and lower needles, so I will hereafter refer to the average trends of all needles. I saw up to 40% of needle water replaced by FUW within the first 24 hours of labelling. According to Berry et al. (2019)’s classification of leaves, the Japanese cedar needles in my study fall in the range of “high foliar water uptake rate” and “high water capacitance” (See Fig. 4 in Berry et al., 2019). Reports on FUW replacement of leaf water vary widely across species and experimental design, with other reports of high foliar water uptake rates including 21% for a temperate conifer (*Araucaria angustifolia*), in overnight fog (See Fig. 3a in Cassana et al., 2016), and nearly 100% in 3 to 8 hours for *Pinus sylvestris* and *Fagus sylvatica* in fog (Lehman et al., 2019; See Lehman et al., 2019 for other examples).

When rain falls on Japanese cedar, the water forms droplets coalesce as they roll down, and are trapped among the bases of needles at the shoot axis, keeping the needles mostly dry (Extended Data Fig. 2, Nanko et al., 2013). This is advantageous for CO_2_ uptake in rainy climates (Hanba et al., 2004), but it limits foliar water uptake from needles, and means that a significant portion “foliar water uptake” may happen via uptake of liquid water through shoot axes instead of vapour through the stomata of needles.

Because plants both absorb and lose water through bidirectional exchange at leaf surfaces (Farquhar & Cernusak 2005; Kim & Lee, 2011; Goldsmith et al., 2017), the drainage water of the above-ground chambers in this experiment included not only run-off from spraying the needles, but also water from inside the saplings (Extended Data Table 3). Without further experiment, it would be hard to say how much heavy FUW from needle water was expelled through leaves back into the environment, but the 45-54% of drainage water composed of natural isotope ratios must have come from inside the saplings or from the soil, while another 4-8% was root-absorbed heavy water.

We can get a sense of the relative rates of foliar water uptake and root water uptake, by noticing that the heavy RUW signals in roots and needles were initially smaller but more persistent than the heavy FUW signal in needles. Fig. 2 is dominated by the heavy FUW signals in needles which spiked dramatically during labelling, but disappeared within four days of the last labelled spray, while heavy RUW signals were more erratic and somewhat less pronounced, but still clearly present in both roots and needles seven days after labelling. I deduce that the turnover of soil water took longer than the turnover of foliar-absorbed precipitation. In other words, foliar water uptake happened at faster rate over a shorter period than root water uptake, creating a lag in the heavy RUW signal. This agrees with published research (Kabeya et al. 2007; Lehman et al., 2018; Lehman et al., 2019). Furthemore, part of the lag can be explained by the time required for root water to be transported to leaves (Ferrio et al. 2018).

The relative size of the isotopic signals of RUW versus FUW in needles can be explained by considering that the drainage water from the ground chambers contained 64-72% natural water (Extended Data Table 3). Even allowing for bidirectional water exchange between roots and soil, I estimate that heavy water available to roots could not have exceeded averaged concentrations of much more than 1/3 during the labelling period. The erratic nature of the heavy water signal may reflect temporal fluctuations of water flux in fine roots and erratic dispersion of water in the soil (Dawson 1993; Treydte et al., 2014) that would make labelling plants through soil water particularly tricky.

### 3.2 Transport of FUW and RUW

The next step is to consider the transport of FUW and RUW within the saplings. The transport of RUW to needles is a direct prediction of the traditional model of transpiration, but what about transport of FUW? Fig. 2 shows that minima of RUW in needle water were accompanied by minima of transpiration, demonstrating that xylem flow changed direction depending on transpiration conditions.

FUW signals in roots were close to zero but still significant in both trees. Cassana et al. (2016) report much larger concentrations of FUW in root water and possible release of FUW through the roots into the soil, creating FUW concentrations in soil+fine root as high as 34%, but this only occurs for water-stressed trees (See Fig. 3 in Cassana et al., 2016). In my experiments, roots were watered only a few minutes after watering the needles, most likely resulting in a bidirectional flow from leaves and roots towards stem, where water potential is lowest (as depicted in Fig. 3B in Schreel & Steppe (2020)). This also explains why drainage water from the ground chambers in this experiment contained only 0.01-0.09% FUW, much too small a signal to detect in the soil. Other authors with better-watered trees also report no detection of FUW in soil (Limm et al., 2009, Berry et al., 2014).

Nevertheless, FUW in roots was first detected on 17 July, the morning after the first labelling, showing the possibility that translocation of phloem sap happened overnight. The first clear RUW signal in needles only appeared later that day. If we refer again to Cassana et al. (2016), we find that direction and volume of advectional flow depend heavily on water potential, and that flow of xylem sap in the direction of leaves to roots is driven by an inverted gradient in the water potential across leaves, stem, and roots. However, maximum FUW in roots observed was 0.5%, which is much smaller than 34% of FUW observed in Cassana et al. (2016). For example, the phloem to xylem flux ratio of water in popular is reported to be 0.19 at night (Windt et al., 2006). Since Cassana et al. (2016) did not find significant amount of FUW in roots of well-watered trees, and flux of water in phloem is much smaller than in xylem, I believe that FUW in roots observed in this study was most likely transported to roots via phloem. Given a phloem translocation velocity of 0.1 m h^-1^ for a conifer (Epron et al., 2019), phloem sap exported from the leaves of saplings like those in this study should be able to reach roots within five hours. In springtime, about 30% of gross primary production is translocated underground via phloem (Epron et al., 2019), and I deduce both active phloem translocation and positive turgor in roots during my experiment because I observed active fine root growth (up to 1 cm per day). Furthermore, phloem thickness is reported to be larger at night when the vapour pressure deficit is lower and phloem water potential is higher (Pfautsch et al., 2015). When trees are well-watered, low osmotic pressure (i.e. low viscosity) of phloem sap enhances active transport of phloem sap (Dannoura et al., 2019). I therefore expect significant active phloem transport under conditions favourable to foliar water uptake. In fact, transport of FUW via phloem is reported in beech leaves (Schreel et al. 2020).

By this reasoning, opening the chambers to outside ventilation and thereby decreasing humidity on 19 July would lead to the normal water potential, increased transpiration, acropetal xylem sap flow, and active water uptake by fine roots the following days. Indeed—except for 22 July, which was a dark day of 100% humidity (data not shown) with a minimum of transpiration—RUW in the needles was observed through 25 July. Absence of RUW in needles on 22 July signifies that sap had been flowing in the inverted, basipetal direction. By then, the trees had been receiving only tap water for three days, and I believe that any heavy FUW still in the needles would have been lost by transpiration or diluted by newly absorbed tap water. However, some of the heavy FUW could still have existed in the phloem of branches and stems, which have capacitance and can store some amount of water (Sevanto et al., 2011, Steppe et al., 2018). I conclude that the combination of high humidity, needle wetting, and minimum transpiration triggered inversion of the water potential gradient and caused heavy FUW in the needles to flow down to branches or stems through xylem, and a small but persistent supply of FUW to the roots through phloem until 25 July.

Conditions on both 24 and 25 July were sunny and dry, and, by the above reasoning, active transpiration would have returned the xylem sap flow the familiar, roots-to-needles direction, replacing tap water in needles with heavy RUW transported from the branch, stem and roots. Heavy RUW in needles reached the second maximum on 25 July at 18:00, but all signals in root and needle water were lost by 6 August, probably due to repeated watering with tap water during and after transplanting and active transpiration under drier weather.

These findings support the hypothesis that the internal dynamics of FUW and RUW are extremely vital to forest hydrology, and help to explain one of the functions of the forest canopy. Anyone who has taken shelter under a tree has experienced canopy interception, and Breshears et al. (2008) have demonstrated that foliar absorption and transportation of intercepted rainfall is sufficient to improve plant water status in *Juniperus monosperma*, which naturally grows in dryland ecosystems. In coniferous forests, the closed canopies keep the forest floor relatively dry at the beginning of rain events.

This is a seasonal effect, with stronger interception during the spring/summer growing season, when the water potential of leaves is more negative (Dunin et al., 1988; Murakami, 2006), and in a mature stand of Japanese cedar, such interception by the canopy can last up to 24 hours from the start of a rain event (See Fig. 5D in Iida et al., 2017). This is similar to the time the trees in my experiment required for needle foliar water uptake to reach equilibrium (roughly estimated as the time between first administration of heavy water and first FUW and RUW signal peaks in Fig. 2).

I believe that foliar water uptake is at least one of the mechanisms driving active canopy interception, and I hypothesize that maximal transport of FUW toward roots occurs at the beginning of rain events, when the canopy interception is greatest (Iida et al., 2017) and the water potential difference between leaves and soil is still large—a variation on the model described by Schreel & Steppe (2020). Even when rain is falling in a forest, the air within the canopy is not saturated (Murakami, 2006, Dunkerley, 2009), suggesting the possibility of canopy-air humidity decrease caused by net foliar “vapour” uptake. Some studies demonstrate “foliar vapour uptake” under high humidity conditions by up-regulation of aquaporin (Laur & Hacke, 2014a; Laur & Hacke, 2014b), even in the absence of physical leaf wetting (Wang et al., 2016; Vesala et al., 2017), which could be tested by future labelling experiments.

### 3.3 Assimilation of FUW and RUW into cellulose

Analysis of the biomass formed during the rainy season of pulse labelling revealed that a significant percentage of hydrogen and oxygen in the cellulose was derived from FUW. This finding literally turns the commonly accepted notion that hydrogen and oxygen of plant biomass derive from water absorbed by roots—as enshrined in textbook figures explaining photosynthesis—upside-down. Although current models (Roden et al. 2000; Farquhar & Cernusak 2005) account for hydrogen and oxygen contribution from atmospheric vapour to hydrogen and oxygen isotopes of tree rings, they do not account for contributions from foliar-uptake liquid water. At least for climates with wet growing seasons, textbook figures explaining water use in photosynthesis are inadequate, and we need new figures that include foliar water uptake (such as Fig. 3). Furthermore, we have discussed evidence that water potential drives transportation of FUW and RUW to set up a FUW/RUW gradient in tissue water. I propose that this FUW/RUW gradient in tissue water is then embedded in the cellulose (as illustrated in Fig. 3). Furthermore, I observed that FUW/RUW gradient extended even within a needle (A. Kagawa, unpublished data). This intra-needle FUW/RUW gradient may give another cause of hydrogen and oxygen isotopic gradient observed within leaf, in addition to Péclet effect (Farquhar & Lloyd, 1993; Farquhar & Gan, 2003).

Let us combine this study’s results with current knowledge to examine how this process might play out. First, as discussed in 3.2, we assume that the uptake and transportation of FUW and RUW create a gradient in the tissue water that can shift according to water status. Photosynthesis assimilates available hydrogen and oxygen from leaf water (and carbon from CO_2_) and embeds the leaf water signal of FUW/RUW into carbohydrates (Epstein et al., 1976; DeNiro & Epstein, 1979; Farquhar et al., 1998). These carbohydrates may be used locally or translocated to other parts of the plant. Although we do not know if and at what rate the carbohydrates exchange oxygen (and hydrogen) with local water during phloem loading, transportation, and unloading (Gessler et al., 2014), even with lack of direct data, we expect carbohydrates dominated by sucrose to carry isotopic signals acquired in production to other parts of the plant, because sucrose has no free carbonyl group to exchange oxygen atoms with local water. However, various authors have found that there is a high rate of exchange during synthesis of cellulose (Hill et al., 1995; Cernusak et al., 2005; Sternberg, 2009), which would encode the local ratio of FUW/RUW into the cellulose.

In leaves, the isotopic signals of biomass should be a function of the integrated isotopic signals of leaf water, dominated by FUW (Barnard et al., 2007; Gessler et al., 2007). Away from the site of photosynthesis, we can expect exchange during cellulose formation to shift the isotopic signals of cellulose from FUW towards RUW as we move in the direction from leaves to roots. Therefore, the signal in tree rings should be a function of the signal in cambial water and depend on the average fractions of carbon-bound hydrogen available for exchange (*q*_ex_) and oxygen available for exchange (*p*_ex_). In my study, the similarity of trends of heavy water distributions in tissues (Fig. 2) and signals in α-cellulose (Extended Data Table 4) in the O/H and H/O tree supported the assumption that *q_ex_*≈*p_ex_* in our approach. The signal in roots should be a function dominated by the RUW signal.

Substitution of *τ*_Ht_ and *τ*_Ot_ with isotope ratios of bulk root biomass (which includes cellulose and other carbohydrates) sampled from 16 July to 25 July gave averaged *pq*_ex_ values of 0.93 for the O/H tree and 0.83 for the H/O tree (Extended Data Table 5), and samples of α-cellulose extracted from roots yielded *pq*_ex_ = 0.90 for the O/H tree and *pq*_ex_ = 1.01 for the H/O tree (calculated from data in Extended Data Table 4 and 5). As time elapsed from the start of labelling, I also observed a gradual increase in *r*, indicating deposition of deuterium and oxygen-18 into newly formed root biomass.

Among reported *pq*_ex_ values, these are on the high end, but not unreasonably so. For various species and growing conditions, other authors have found that, *q*_ex_ during biomass synthesis to be close to 0.40 (Yakir & Deniro, 1990; Terwilliger & Deniro, 1995; Roden et al., 2000; Waterhouse et al., 2002) and *p*_ex_ usually between 0.40 and 0.50 (Sternberg & Deniro, 1983; Sternberg et al., 1986; Yakir & Deniro, 1990; Hill et al., 1995; Roden et al., 2000; Helliker & Ehleringer, 2002; Sternberg et al., 2003; Cernusak et al., 2005; Sternberg et al., 2006), but as low as 0.32 (Roden et al., 2000). Under certain circumstances, however, *p*_ex_ can be much higher. For example, *p*_ex_ is known to show seasonal variations between spring and summer, with the highest exchange observed at the beginning of the growing season (*p*_ex_ = 0.76 for *Fagus sylvatica*, Offermann et al., 2011), when elevated temperatures are known to promote enzymatic conversion of starch to sugar to supply energy for new cell wall formation (Begum et al., 2010). Variable *p*_ex_ from 0.40 to 1.0 have been reported (Schmidt et al., 2001; Sternberg et al., 2006) with especially high *p*_ex_ values under water stressed and high salinity conditions (Ellsworth & Sternberg, 2014). For a supply of assimilate to be maintained, phloem transport has to stay unaffected by water stress and high salinity, and, for example, *Ricinus communis* under water stress increases sucrose content in the sieve tubes by osmoregulation (Smith & Milburn, 1980). Incidentally, Song et al. (2014) also studied *Ricinus communis* to find positive correlation between *p*_ex_ and the turnover time of carbohydrate (sucrose) pool available for cellulose synthesis.

Since the trees on my study were well–watered, one potential explanation for high *p*_ex_ (and *q*_ex_) could be that foliar water uptake caused leaf water and the carbohydrates contained within to be transported to the stem and roots via phloem (Laur & Hacke 2014a; Schreel et al. 2020), thereby increasing phloem sugar content―and hence turnover time of carbohydrate pool― at the branch, stem and roots. As cells at the cambial zone absorb water and expand, solutes in their cytoplasm are diluted. To maintain constant osmotic pressure in cytoplasm, expanding cells actively break down sucrose into glucose and fructose (Koch 2004), which may explain the unusually high exchange of oxygen (and hydrogen) between sugars and local water that I observed. Song et al. (2014) hypothesized that *p*_ex_ and the turnover time of the carbohydrate (sucrose) pool is positively related. This hypothesis could be tested with further labelling experiments.

Eventually, the labelling signal of water inside trees would be either incorporated into the structural components, such as tree-ring cellulose, or released back into the environment. Although I found significant labelling signals in new roots that were sampled during the labelling period (Extended Data Table 4 and 5), replanting the trees on July 26 also allowed me to later isolate roots formed in the post-labelling period, when there would have been less, if any, labelled primary photosynthate available for cellulose production. When I took root samples the following January, I did not find significant labelling signals in any of the thick, post-labelling roots (diameter up to 3 mm) sampled from all six labelled trees (*P* = 0.095, Extended Data Table 6). In a previous study, my laboratory successfully detected signals from photosynthate labelled with ^13^CO_2_ in July in roots formed later that year (Kagawa et al., 2006b). Since I found no such D and ^18^O signals in the present study, I deduce a complete exchange of D and ^18^O signals in the carbohydrate pool with medium water and/or fast turnover of the carbohydrate pool within the six months between labelling in July and sampling the following January. Since 83-93% of hydrogen and oxygen in substrates for roots was replaced with root water within a week (Extended Data Table 5), hydrogen and oxygen of the carbohydrate pool might have been turning over at a much faster rate than carbon. In the coming years, I plan to analyse hydrogen and oxygen isotope ratios of tree rings formed in years following labelling to see if hydrogen and oxygen isotope signals also appear in later rings, as ^13^C signals appear in later rings after ^13^CO_2_ labelling (Kagawa et al. 2006a).

Current mechanistic models explaining tree-ring hydrogen and oxygen isotope ratios are based mainly on experiments in which trees were grown in hydroponic facilities without leaf wetting (Roden et al., 2000). However, in the present study, more than half of the hydrogen and oxygen in cellulose extracted from the current-year’s branch wood was derived from FUW, so this practice is clearly inadequate. We can make a comparison to carbon allocation patterns of trees, which can be explained by a source/sink model (Lacointe, 2000). There is only one source of carbon for autotrophic trees: Leaves (to be more exact, photosynthetic organs). However, there are at least two sources of hydrogen and oxygen: Foliar-uptake water and root-uptake water. According to source-sink models of carbon allocation, each non-green organ (sink) draws carbon from the nearest source with the shortest distance. Similarly, in this study, cellulose of leaves contained mostly foliar uptake water, roots contained mostly root-uptake water, and branch wood contained roughly equal parts of foliar-uptake water and root-uptake water (Extended Data Table 4), thus suggesting that this rule also holds true for hydrogen and oxygen allocation of trees during rain events. Because foliar uptake of liquid water was accompanied by an increased exchange of hydrogen and oxygen (up to 100%) between local water and carbohydrate pool (Extended Data Table 5), I conjecture that, during a rain event following days of sunshine, hydrogen and oxygen isotopic information in sugars, which carries leaf evaporative enrichment signals (elevated δD and δ^18^O) from the time of photosynthesis, may be mostly overwritten by local water at the cambium. A significant portion of hydrogen and oxygen incorporated into tree rings formed during rainy season should therefore derive directly from foliar-uptake water, especially for trees in humid regions, such as Monsoon Asia. Perhaps this can explain some recently published findings such as unusual δD and δ^18^O values in the earlywood of Japanese oak (Nabeshima et al., 2018).

Foliar water uptake becomes most active when water potential difference between leaves and roots is largest (i.e. when leaves are wet and roots are dry; Eller et al. 2013; Cassana et al. 2016). Such conditions may happen during a rain event at a mature forest with closed canopy, where larger interception at the canopy and lesser infiltration into the soil are expected. Among forests in the world, coniferous forests with closed canopy shows the highest canopy interception loss, up to 48% of total rainfall (Hörmann et al. 1996). Indeed, high coherence in tree-ring δ^18^O is reported in such forests (Roden et al., 2008; Roden et al., 2009; Roden et al., 2011). On the contrary, I expect a low coherence in arid regions, where leaves infrequently get wet and trees rely more on ground water (Qin et al. 2015). On one hand, future studies may find that the percentage of FUW in stem wood turns out to be somewhat lower than the percentages in branch wood where samples were taken in this study. However, since the latter half of earlywood formation of Japanese trees overlaps with the rainy season, and radial growth reaches maximum speeds during this period (Kagawa et al., 2005; Rossi et al., 2006), I expect that foliar-uptake liquid water to make a significant contribution to tree-ring hydrogen and oxygen isotopes in Monsoon Asia. If this is correct, current mechanistic models explaining hydrogen and oxygen isotope ratios of leaf water and tree rings will need to be revised to account for contributions from foliar water uptake.

## 4 Conclusions

As far as I know, this is the first study to quantify relative contributions of foliar-uptake water and root-uptake water to the biomass of terrestrial plants. I found that foliar water uptake reached equilibrium within 12-24 hours in *Cryptomeria japonica* saplings, and the ratio of foliar-uptake water and root-uptake water increased from roots to branches to leaves, creating isotopic patterns that were embedded in cellulose. More than half of the hydrogen and oxygen in new branch wood originated from foliar-uptake liquid and vapour water. Although these results may be most representative of trees with rainy growing seasons, they clearly demonstrate the need to overhaul the commonly accepted view that hydrogen and oxygen in plant biomass are mainly supplied from soil water. However, the results of these experiments cannot be simply extrapolated to mature trees, because mature trees may rely more on stored carbohydrates remobilized from parenchyma cells, and water stored in heartwood may move radially to cambium (Nakada et al. 2019) and may be eventually incorporated into biomass. Future research on mature trees over a longer time period is still required to determine, exactly how much hydrogen and oxygen in plant biomass originates from foliar-uptake water.

The dual-isotope pulse-labelling method can be used to further explore this topic and other biological processes related to this study, such as transport processes of foliar-uptake water, and clarification of whether foliar-uptake water is transported via xylem, phloem, or some combination of the two. We also need to test whether net foliar water uptake of trees can happen from near-saturated air even in the absence of physical leaf wetting, and if so, at what humidity (Yan et al., 2015). We might also address how much foliar-uptake water is used for production of molecular oxygen through photosynthesis. In the future, experiments with stable isotopes, if conducted in a closed system with closely monitored isotopic mass balances between plants and the surrounding environment, may even enable us to quantify the absolute net foliar uptake of water into plants. Another interesting subject would be application to isotope dendroclimatology. For example, I would hypothesize that tree-ring cores sampled from upper and lower parts of the stem may have isotopic differences that are most obvious between wood increments formed during rain events, because the upper core may rely more on foliar-uptake water. In conclusion, the dual labelling method with HDO and H_2_^18^O first implemented in this study offers a new tool to study the uptake, transport, and assimilation processes of water in terrestrial plants, and successfully enabled independent labelling of foliar-uptake water and root-uptake water. Using this method, I found conclusive evidence that foliar-uptake liquid water can be a significant source of hydrogen and oxygen in plant biomass, such as tree rings, formed during rainy season.

## M Methods

### M.1 Sample trees and labelling experiment

For this experiment, I used eight two-year old saplings of Japanese cedar, grown at the Forestry and Forest Products Research Institute in Tsukuba, Japan. Each sapling grew in a pot (inner diameter 45 mm, depth 135 mm), filled with coconut coir and fertilized with slow-release nitrogen, phosphorus, and potassium. The pots were moved to a nearby greenhouse for acclimatization one month before the labelling experiment. During this period, they were watered every 2-3 days to field capacity with tap water. The greenhouse was well ventilated with open side windows, and equipped with a glass roof to shelter the saplings from rain. Average sapling height was 45 ± 14 cm at the time of labelling.

The saplings were divided into three groups: The O/H group consisting of trees A, B, and C would have their aboveground parts supplied with heavy oxygen water and their belowground parts supplied with heavy hydrogen water. The H/O group consisting of trees D, E, and F would have their aboveground parts supplied with heavy hydrogen water and their belowground parts supplied with heavy oxygen water. The control group consisting of two trees G and H would be supplied only tap water.

On July 16 at 15:00, I installed separate chambers constructed from transparent, resealable plastic bags around the aboveground parts and belowground parts of each sapling, as shown in Fig. 1a. I reinforced the seal around the stem where it exits the aboveground chambers with Gaffer tape, leaving a 5-cm gap between chambers, where I gently wrapped each stem in paper towel. I turned the bottom hem of each aboveground chamber up towards the inside to create a gutter leading condensation towards a drainage bottle. Soil drained to a bottom corner in the ground chambers. Throughout the experiment, the air inside each aboveground chamber was gently circulated by an electric fan. The two control saplings were placed immediately downwind of the experimental saplings to detect potential interference of heavy-water vapour escaping the chambers. To prevent condensation, chamber seals were left open until after the first labelling procedure.

On 16 July at 18:00, around sunset, I used a sprayer to wet the surfaces of the upper needles of tree A with heavy oxygen water (0.2 atom% ^18^O). After closing the top seal, I opened the side seal and similarly wetted the middle and lower needles, injecting a total of 15.4 ml of heavy oxygen water into the aboveground chamber. I then closed the side seal, leaving open a small vent just large enough for atmospheric gases to enter the chamber and maintain a CO_2_ concentration above 200 ppm. Next I opened the seal of the ground chamber. I used a pipette to uniformly wet the upper surface of the soil with 12 ml of heavy hydrogen water (1.1 atom% D), then closed the seal, except for a CO_2_ vent (Fig. 1a). I repeated the same procedure on trees B and C, and repeated the whole process for the three saplings in the H/O group, but with the heavy water supplies swapped for the aboveground and belowground parts. All of this was completed within 30 minutes of 18:00. On July 17 and 18 at 06:00, 12:00, and 18:00, I repeated this labelling process for both groups of trees. In total, the aboveground parts of each tree received 108 ml of heavy water and the belowground parts of each tree received 84 ml of heavy water. I chose these values in consideration of reports that, under the trees’ typical growing conditions, 22 % of rainfall does not reach the ground due to interception mechanisms including foliar water uptake (Murakami, 2006; Breshears et al., 2008).

After the last administration of heavy water, I kept the aboveground chambers sealed except for CO_2_ vents for 24 hours to emulate high-humidity conditions for one day following a rain event. On 19 July at 18:00, I collected the drainage water and opened the chambers to outside ventilation (Fig. 1b) until 20 July at 06:00, when I removed the chambers completely. I weighed the saplings on 20, 21, 22, 24, and 26 July to measure transpiration (Seiler & Johnson, 1988), and watered them with tap water (15.4 ml to aboveground parts and 12 ml to belowground three times a day) through 18:00 on 26 July, at which point I transplanted saplings to 4 litre pots to accommodate growing root systems and watered the pots to field capacity three times. From this date onward, I watered saplings with tap water every 2-3 days to field capacity during the growing season (until Oct. 2019), including wetting the needles and watering the soil, then every 7 days during the dormant period (after Nov. 2019). I could observe moderate shoot and root growth between August and October 2019. Temperature and CO_2_ concentration were monitored inside the chambers from 16 to 19 July, and outside the chambers from 15 July to 7 August.

I sampled needles from upper, middle and lower branches, and new roots (easily distinguishable from older roots by their white colour) periodically from 16 July at 17:30 (before labelling) through 25 July (6 days after the last labelling) and sampled needles again on 28 July and 6 August. I limited samples to volumes the saplings could tolerate without damage and reduced sampling frequency as necessary, as shown in Fig. 2. When samples were wet, I wiped wet surfaces dry, then sealed all samples in air-tight vials and put them immediately into a freezer (−14 °C) located at the greenhouse before transfer to a larger freezer (−30 °C) closer to our microbalance.

The following January, I sampled roots and current-year branches for α-cellulose extraction. The branch was sampled halfway between the positions where I sampled upper and middle needles. To minimize damage and keep the trees healthy for later observation, I used roots formed within the soil added around the original coconut coir. I extracted α-cellulose from these samples and also from needles and roots sampled previously on 25 July according to the methods of Laumer et al. (2009) and Kagawa et al. (2015). Needles, phloem, and epidermis were removed from current-year branches. Needles were dried, ground in a steel ball mill (ICL, Wig-L-Bug Model 30) and extracted with 3:1 chloroform/ethanol. Phloem+epidermis and xylem samples were sliced by razor into thin slivers. Chloroform extracted needles, slivered phloem+epidermis, and slivered xylem samples were next extracted with ethanol (60◻, 48 h) and reverse osmosis water (100◻, 48 h), acidified sodium chlorite (70 ◻, 12 h), and NaOH (80 ◻, 12 h). Finally, the resulting α-cellulose was rinsed, ultrasonically homogenized, and freeze-dried (Laumer et al. 2009).

### M.2 Stable isotope sample preparation and analysis

For this analysis, in addition to measurements of the control trees G and H, I measured the isotope ratios in the water of upper, middle, and lower needles and fine roots of one randomly selected O/H sapling (tree C) and one randomly selected H/O sapling (tree F); the organic matter of upper, middle, and lower needles, and fine roots of one O/H sapling (tree C) and one H/O sapling (tree F); the cellulose of needles, phloem+epidermis, xylem, and roots of all saplings; the drain water from the aboveground and ground chambers of all saplings; and the heavy hydrogen water, heavy oxygen water, and local (Tsukuba) tap water.

Before analysis of each plant sample, I removed the vial from the freezer and let it approach room temperature for a few minutes to prevent condensation. Then, working quickly to minimize evaporative losses, I removed ca. 1-mm segment from the cut surfaces of each needle or piece of fine roots to prevent potential contamination from heavy water absorbed from the cut surfaces, then divided the rest of each piece into three segments of roughly equal length and placed the central segments in preweighed smooth-walled tin capsules designed for liquids. I folded over the lip of each capsule and pressed it closed three-times to make the seal airtight. I weighed these capsules, placed them in the carousel of a Zero Blank autosampler (Costech), and ran the rotary vacuum pump connected to the autosampler for more than one hour before weighing the capsules again to check for leaks. Meanwhile, I placed the remaining outer pairs of needle and root segments in preweighed tin capsules designed for solids, weighed them, dried them in a vacuum oven at 70 ◻ for more than 24 hours, and weighed them again to calculate the water content of needles and root segments. I also prepared samples of heavy waters, drainage water from the chamber vials, and reference water standards in smooth-walled tin capsules. For reference standards, I used artificially enriched and natural water (IAEA-607, IAEA-608, and IAEA-609, International Atomic Energy Agency, our internal standard) and natural cellulose (IAEA-C3 cellulose, Merck cellulose) with known isotope ratios measured against the VSMOW standard (detailed in the next section). To increase precision, I prepared five or more capsules from each sample of water and cellulose, except when limited by sample volume (Olsen et al. 2006), and two to five capsules from each needle and root sample, depending on the amount of samples available. However, I successfully analysed only one out of two capsules of roots of O/H sapling sampled at 06:00 and 18:00 on 18 July, and at 18:00 on 19 July, due to improper technique (Fig. 2a).

To analyse the hydrogen and oxygen isotope ratios of solid, liquid and wet tissue (= solid+liquid) samples, I used a high-temperature pyrolysis system (Hekatech, HTO) coupled via thermal conductivity detector to a mass spectrometer (Thermo-Finnigan, Delta V Advantage) with a longer gas chromatography column (molecular sieve 5A, 2 m, i.d. 5 mm) to achieve clear separation of hydrogen, nitrogen, and carbon monoxide gas (Brand et al., 2009). To minimize background moisture and nitrogen gas from the samples, I opened the three-way valve on the autosampler to the vacuum pump for 30 secs, then switched the valve to the helium gas supply for 30 seconds, and repeated this procedure ten or more times until the background levels (mass=28, 29, 30) stabilized, before initiating analysis (Kagawa et al., 2015). To minimize the impact of sample-to-sample memory, I grouped together plant and water samples with similar isotope ratios in the analytical sequence. I replaced the pyrolysis tube as necessary due to tin accumulation, typically every 50 samples for smooth-walled tin capsules.

### M.3 Calculation of isotope data

I based calculations of excess D and ^18^O of labelled samples on published methodologies for calculating excess ^13^C of ^13^CO_2_ pulse-labelled samples as follows (Boutton, 1991, Kagawa et al., 2006b). For each sample, δD and δ^18^O (δ) were first converted to absolute isotope ratio (*R*_sample_):

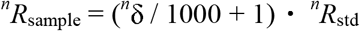

where *n* is the mass number of the hydrogen or oxygen isotopes and *R*_std_ is the absolute isotope ratios of Vienna Standard Mean Ocean Water (VSMOW: ^2^*R*_std_ = 0.00015576, ^17^*R*_std_ = 0.000379, and ^18^*R*_std_ = 0.0020052; De Wit et al., 1980; IAEA 2006). The D and ^18^O abundance ratios (*A*) were calculated as follows:

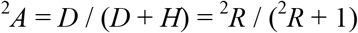

and

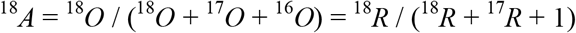

Then excess D and ^18^O abundance (*∆^n^A*) was calculated as the deviation from natural isotopic values:

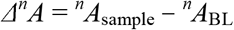

where *A*_BL_ and *A*_sample_ are abundance ratios of the samples taken before and after labelling, respectively. Excess abundance of tissue water (*∆A*_TW_) was calculated from isotopic mass balance as,

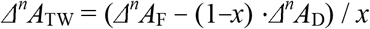

where *∆A*_F_ and *∆A*_D_ are the excess abundances of fresh and dry samples of needles or roots, and *x* refers to the portion of hydrogen or oxygen contributed by tissue water (leaf water or root water) to the total hydrogen or oxygen of the fresh sample, calculated from the water and elemental content of fresh and dry samples. (See Gan et al. (2004) for details.) I then introduced *τ*H and *τ*O, the percentages of hydrogen and oxygen, respectively, in samples replaced by atoms from heavy water and calculated as,

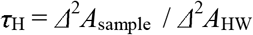

and

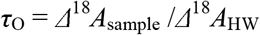

where *∆^n^A*_HW_ is the excess abundance of heavy water compared to local tap water.

Hydrogen and oxygen isotope fractionations show similar patterns within hydrological cycles along the meteoric water line (Dansgaard 1964), but they show difference within biological systems, at the leaf water level (Yakir et al. 1990), at the cambium (Yakir and DeNiro, 1990), and at tree rings (Nabeshima et al. 2018, Nakatsuka et al. 2020). Such deviations in isotopic fractionation between hydrogen and oxygen seem to be related to metabolic activities (Lehmann et al. 2020, Nakatsuka 2020), and one possible cause for such deviations is respiration, because oxygen is eliminated by both carbon dioxide and water, and hydrogen would be eliminated only by the water. In fact, deviations between oxygen and hydrogen isotopes are used to assess amount of respired CO_2_ in humans and animals (Schoeller & Santen 1982, Speakman 1997), and similar phenomenon might be happening in plants. To offset any effect of the chemical difference between hydrogen and oxygen (Newberry et al. 2017), and to calculate the relative contribution of FUW and RUW assimilated into biomass (I focus on cellulose in this paper) by weight, I calculated *τ*_Cell_, the weight percentage of hydrogen+oxygen in cellulose replaced by heavy water. Since the cellulose samples were not nitrified, I assumed hydrogen atoms in the three OH bonds of each cellulose unit were replaced by natural hydrogen atoms while boiling in reverse osmosis water overnight, because OD (oxygen-deuterium) moieties of cellulose can be easily rehydrogenated by soaking cellulose in water at 25 °C (Horikawa & Sugiyama, 2008), and therefore multiplied cellulose *τ*_H_ by a factor of 1.43 (=10/7).

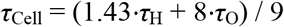

To check for differences between O/H and H/O trees, I compared the corresponding *τ* of α-cellulose across four different parts (needles, phloem+epidermis, xylem and roots, Extended Data Table 4), using an *F*-test (two-way ANOVA with three replication, MS Excel 2016).

Because hydrogen and oxygen show similar biochemical behaviour within the various parts of trees, hydrogen and oxygen isotope ratios of organic matter of various tissues (δD_t_ and δ^18^O_t_)―including needles, phloem, epidermis, xylem, and roots―can be explained by the same mechanistic model as follows (Roden et al., 2000):

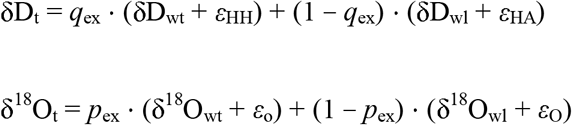

where the subscripts wt refers to local tissue water at non-green parts of the tree and wl refer to leaf water at the chloroplast. The hydrogen and oxygen available for exchange are represented by *q_ex_* and *p_ex_*, respectively. The hydrogen isotope fractionation factors associated with heterotrophic and autotrophic metabolism are expressed as *ε*_HH_ (= +158 ‰) and *ε*_HA_ (= −171 ‰), respectively. For oxygen, there is no need to distinguish between the two, hence both are expressed as *ε*_O_ (= +27 ‰) (Roden et al. 2000).

We are specifically interested in δD_t_ and δ^18^O_t_ of new tissues. *∆*D_t_ and *∆*^18^O_t_ can be related to *τ*_H_ and *τ*_O_ of whole tissue sample (*τ*_Ht_ and *τ*_Ot_), as follows:

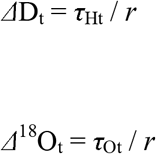

where *τ*_Ht_ and *τ*_Ot_ can be calculated from measured isotope ratios by mass spectrometer, and *r* is proportion of H and O in new tissue to those of whole sampled tissue.

### M.4 Calculation of isotope exchange

Compared to oxygen, less research is available concerning hydrogen isotope exchange (Yakir & DeNiro, 1990; Terwilliger & DeNiro, 1995; Roden & Ehleringer, 1999b; Waterhouse et al., 2002), simply because it is more difficult to analyse natural-abundance isotope ratios of carbon-bound hydrogen in plant biomass. I assumed that *q_ex_*≈*p_ex_* (an assumption justified in the Discussion section, 3.3.) in order to simplify calculations. (Roden & Ehleringer, 1999b, Roden et al. 2000, Yakir & DeNiro 1990). Considering that the excess δD and δ^18^O of the heavy water (Extended Data Table 2) were more than 400 times larger than the absolute values of *ε*_HH_, *ε*_HA_, and *ε*_O_, I deemed contributions from *ε*_HH_, *ε*_HA_, and εO to be negligible compared to δD_wt_, δD_wl_, δ^18^O_wt_, and δ^18^O_wl_. Therefore, to estimate fractions of hydrogen and oxygen exchange with the water at the cambium, I used the following formulas:

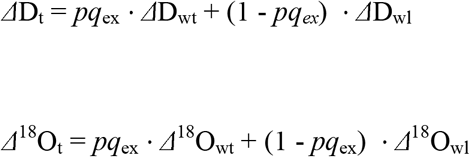

where *∆*D_wt_ and *∆*^18^O_wt_ are *τ*_H_ and *τ*_O_ of tissue water and ∆D_wl_ and *∆*^18^O_wl_ are *τ*_H_ and *τ*_O_ of leaf water. *∆*D_wl_, *∆*^18^O_wl_, *∆*D_wt_, and *∆*^18^O_wt_ can be calculated as time-integrals of *τ*_H_ and *τ*_O_ values of leaf water and root water from data in Fig. 2 (Extended Data Table 5). By solving these equations simultaneously for *r* and *pq*_ex_, fractions of hydrogen and oxygen exchange can be calculated from measured isotopic data as follows.

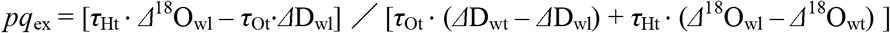

I examined roots in particular because roots were the only non-green tissue I could repeatedly sample without damaging trees.

## Acknowledgements

The author thanks Gerd Helle, and Masaki Sano for development of the research ideas, Kiyosada Kawai, Tayoko Kubota, Kazuki Nanko, and Shin’ichi Iida for comments, and Yuji Kominami for experimental support. Jennifer Lue revised the first draft of this paper. This work was supported by JSPS KAKENHI Grant Number JP19K06179.

## Author Contributions

AK has done all the work.

## Competing financial interests

None declared.

## Data availability

All data supporting the findings of this study are available within the paper. Any additional information may be obtained from the corresponding author upon reasonable request.

## Extended Data Tables

**Extended Data Table 1.**
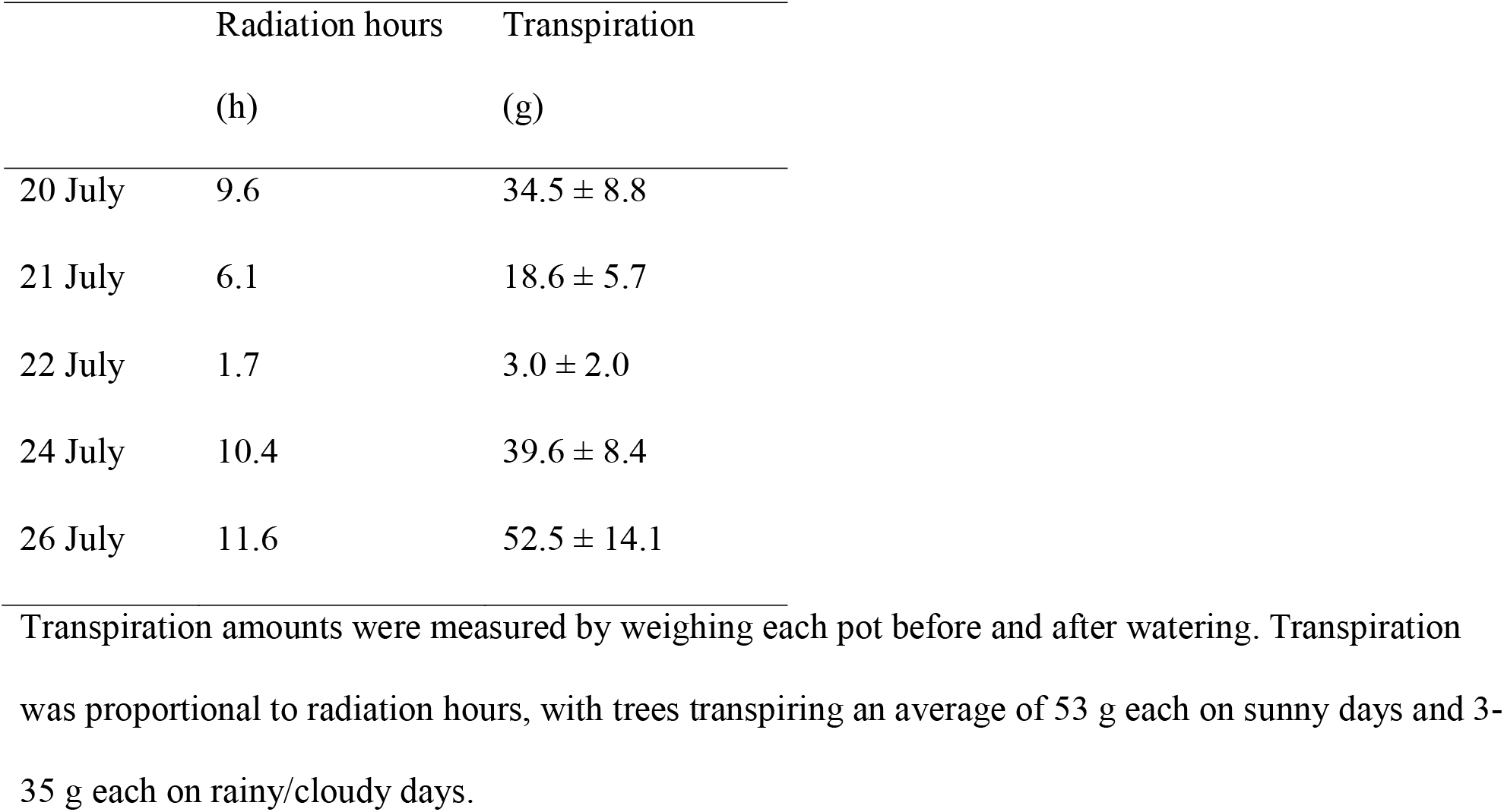
Hours of solar radiation and amounts of transpiration.

|  | Radiation hours | Transpiration |
| --- | --- | --- |
|  | (h) | (g) |
| 20 July | 9.6 | $34.5 \pm 8.8$ |
| 21 July | 6.1 | $18.6 \pm 5.7$ |
| 22 July | 1.7 | $3.0 \pm 2.0$ |
| 24 July | 10.4 | $39.6 \pm 8.4$ |
| 26 July | 11.6 | $52.5 \pm 14.1$ |

Transpiration amounts were measured by weighing each pot before and after watering. Transpiration was proportional to radiation hours, with trees transpiring an average of 53 g each on sunny days and 3-35 g each on rainy/cloudy days.

**Extended Data Table 2.**
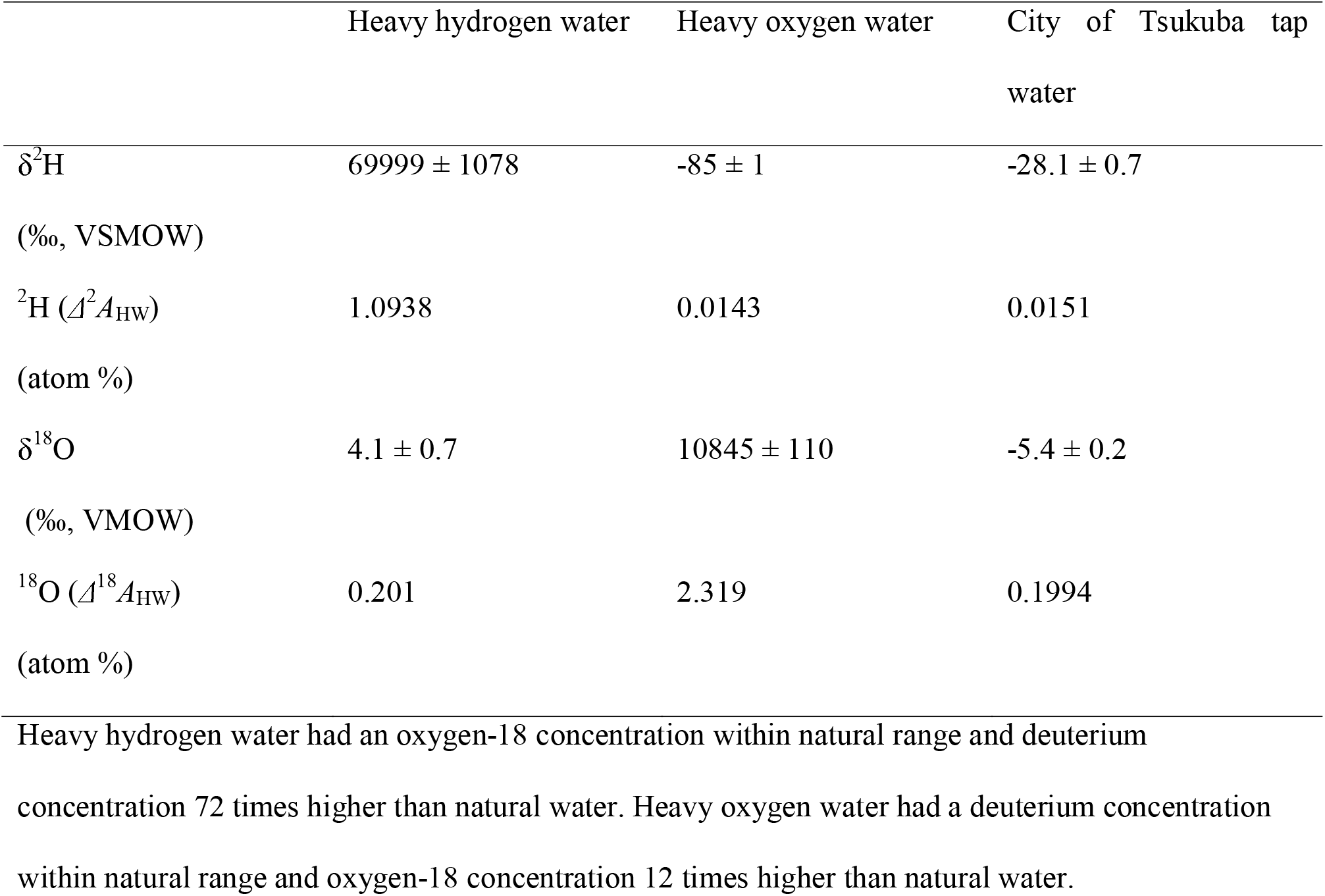
Isotope ratios and atom abundances in heavy water and tap water.

**Extended Data Table 3.**
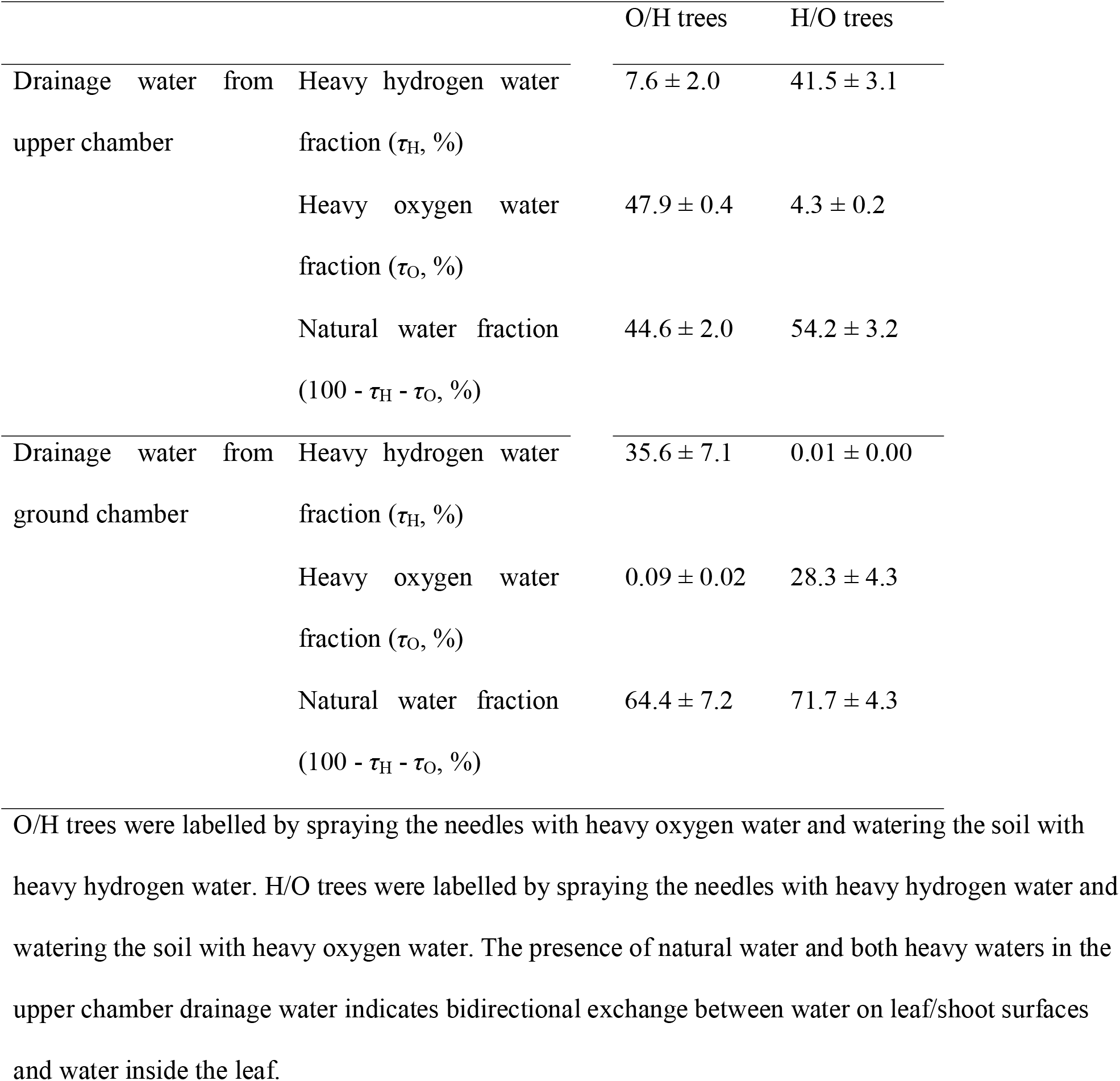
Proportions of heavy water detected in drainage water.

**Extended Data Table 4.** Origins of hydrogen and oxygen in α-cellulose.

|  | Foliar-uptake<br>water<br>(FUW, %) | Root-uptake<br>water<br>(RUW, %) | Foliar-uptake water / root-uptake<br>water<br>(FUW/RUW) |
| --- | --- | --- | --- |
| Average of O/H and H/O trees | ($\tau_{\text{Cell}}$) | ($\tau_{\text{Cell}}$) | ($\tau_{\text{Cell}}/\tau_{\text{Cell}}$) |
| Needles | $72 \pm 2$ | $28 \pm 4$ | $2.2 \pm 0.1$ |
| Phloem+epidermis | $73 \pm 4$ | $26 \pm 5$ | $2.3 \pm 0.2$ |
| Xylem | $57 \pm 4$ | $46 \pm 3$ | $1.2 \pm 0.1$ |
| Roots | $19 \pm 7$ | $77 \pm 7$ | $0.2 \pm 0.4$ |
| O/H trees | ($\tau_{\text{O}}$) | ($\tau_{\text{H}}$) | ($\tau_{\text{O}}/\tau_{\text{H}}$) |
| Needles | $72 \pm 2$ | $28 \pm 2$ | $2.6 \pm 0.1$ |
| Phloem+epidermis | $74 \pm 5$ | $26 \pm 5$ | $2.9 \pm 0.2$ |
| Xylem | $58 \pm 4$ | $42 \pm 4$ | $1.4 \pm 0.1$ |
| Roots | $18 \pm 7$ | $82 \pm 7$ | $0.2 \pm 0.4$ |
| H/O trees | ($\tau_{\text{H}}$) | ($\tau_{\text{O}}$) | ($\tau_{\text{H}}/\tau_{\text{O}}$) |
| Needles | $67 \pm 4$ | $33 \pm 4$ | $2.1 \pm 0.1$ |
| Phloem+epidermis | $68 \pm 5$ | $32 \pm 5$ | $2.1 \pm 0.2$ |
| Xylem | $53 \pm 3$ | $47 \pm 3$ | $1.1 \pm 0.1$ |
| Roots | $24 \pm 8$ | $76 \pm 8$ | $0.3 \pm 0.4$ |

$\tau_{\text{H}}$ is the percentage of hydrogen atoms in α-cellulose replaced by heavy hydrogen water. $\tau_{\text{O}}$ is the percentage of oxygen atoms in α-cellulose replaced by heavy oxygen water. $\tau_{\text{Cell}}$ is weight proportion of (hydrogen+oxygen) in cellulose replaced by heavy water. Most of the hydrogen and oxygen incorporated into new leaf cellulose was derived from FUW, while that of new root cellulose was derived mostly from RUW, and that of new branch xylem was derived from roughly equal parts of FUW and RUW.

**Extended Data Table 5.** Proportion of H and O exchange between root water and root biomass.

|  |  | Root sampling date |  |  |  |  |  |  |  |  |
| --- | --- | --- | --- | --- | --- | --- | --- | --- | --- | --- |
| Tree | Variable | 7/17 6h | 7/17 18 h | 7/18 6h | 7/19 18h | 7/20 18h | 7/21 18h | 7/22 18h | 7/25 18h | Average |
| O/H | $pq_{ex}$ | **0.98** | **0.95** | **1.00** | **0.98** | **0.90** | **0.90** | **0.91** | **0.79** | **0.93** |
| tree | $r$ | 0.04 | 0.06 | 0.18 | 0.25 | 0.43 | 0.44 | 0.44 | 0.37 | 0.28 |
| | $\tau_{Ht}$ | 0.44 | 0.76 | 2.31 | 3.41 | 5.31 | 5.42 | 5.31 | 3.32 | |
| | $\Delta D_{wl}$ | 0.00 | 0.51 | 1.30 | 3.33 | 3.96 | 3.61 | 3.11 | 2.72 | |
| | $\Delta D_{wt}$ | 10.36 | 14.23 | 13.00 | 13.78 | 13.23 | 13.43 | 12.86 | 10.61 | |
| | $\tau_{Ot}$ | 0.02 | 0.08 | 0.04 | 0.23 | 1.10 | 0.96 | 0.74 | 0.87 | |
| | $\Delta^{18}O_{wl}$ | 19.25 | 28.33 | 30.44 | 28.70 | 23.67 | 19.17 | 16.00 | 10.71 | |
| | $\Delta^{18}O_{wt}$ | 0.10 | 0.15 | 0.18 | 0.28 | 0.27 | 0.25 | 0.22 | 0.15 | |
| H/O | $pq_{ex}$ | **0.57** | **0.71** | **0.83** | **0.80** | **0.92** | **0.85** | **0.86** | **1.08** | **0.83** |
| tree | $r$ | 0.07 | 0.07 | 0.21 | 0.29 | 0.43 | 0.45 | 0.46 | 0.30 | 0.29 |
| | $\tau_{Ht}$ | 2.92 | 1.53 | 2.42 | 3.05 | 1.43 | 1.74 | 1.12 | 0.94 | |
| | $\Delta D_{wl}$ | 12.74 | 22.53 | 26.29 | 24.46 | 20.04 | 16.11 | 13.50 | 9.12 | |
| | $\Delta D_{wt}$ | 1.06 | 1.08 | 0.85 | 0.66 | 0.59 | 0.65 | 1.10 | 1.67 | |
| | $\tau_{Ot}$ | 0.78 | 0.98 | 3.55 | 4.47 | 5.68 | 4.39 | 2.63 | 5.59 | |
| | $\Delta^{18}O_{wl}$ | 0.00 | 0.01 | 0.19 | 0.86 | 1.35 | 1.93 | 2.10 | 1.60 | |
| | $\Delta^{18}O_{wt}$ | 2.82 | 6.52 | 9.02 | 9.79 | 9.00 | 8.23 | 7.46 | 5.89 | |

**Extended Data Table 6.** Isotope ratios of α-cellulose extracted from roots sampled in January.

| | $\delta D$ (‰ VSMOW) | $\delta^{18}O$ (‰ VSMOW) |
| --- | --- | --- |
| O/H trees | $-65.7 \pm 3.4$ | $34.0 \pm 0.1$ |
| H/O trees | $-42.5 \pm 3.8$ | $35.3 \pm 0.3$ |
| Control trees | $-10.2 \pm 4.0$ | $33.8 \pm 0.5$ |

Roots for this analysis were sampled in January, 2020, approximately six months after labelling. No significant increase of D and $^{18}O$ signals were found, indicating a complete exchange of D and $^{18}O$ in the carbohydrate pool with medium water and/or fast turnover of the carbohydrate pool within the six months

## Extended Data Figures

**Extended Data Fig. 1.**
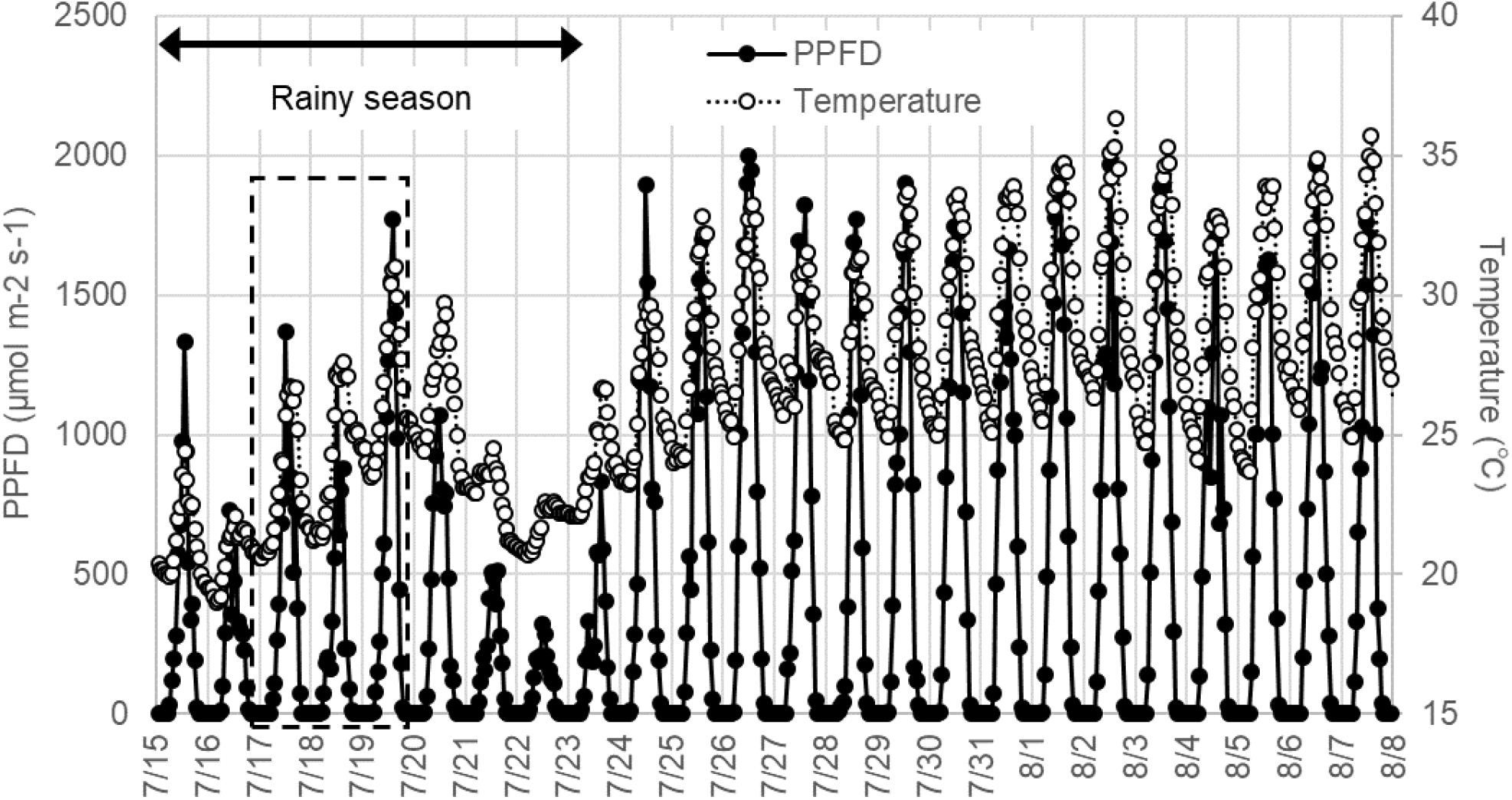
Hourly temperature and PPFD data. Labelling with heavy water was conducted from 16 July at 18:00 to 19 July at 18:00 (dashed box). Photosynthetic photon flux density (PPFD) was estimated from radiation data.

**Extended Data Fig. 2.**
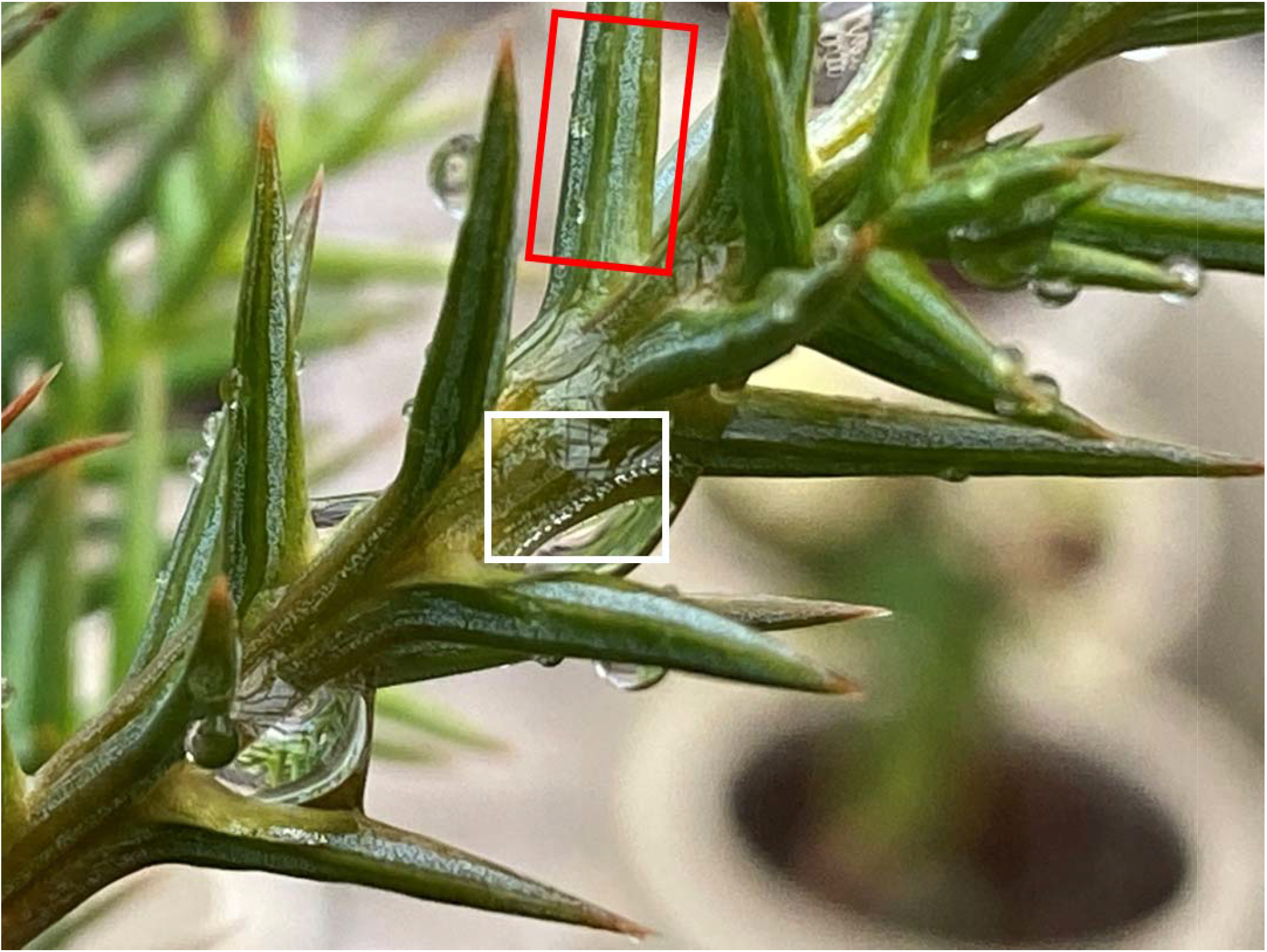
Droplets on *Cryptomeria japonica* needles and shoots. Because of the low-wettability of Japanese cedar needles, needles were mostly free from water droplets, so stomatal bands could function unimpeded by water (red box). Droplets tended to roll down needle surfaces to form larger drop with longer residence time at the shoot axis surface between needle bases (white box).

## Notes

### Competing Interest Statement

The authors have declared no competing interest.

### Summary of Updates

Manuscript was revised based on the personal communications with the peers.

## References

Azuma, W., Ishii, H. R., Kuroda, K. & Kuroda, K. Function and structure of leaves contributing to increasing water storage with height in the tallest *Cryptomeria japonica* trees of Japan. Trees 30, 141–152 (2016).

Azuma, W. A., Nakashima, S., Yamakita, E., & Ohta, T. Water adsorption to leaves of tall *Cryptomeria japonica* tree analyzed by infrared spectroscopy under relative humidity control. Plants 9, 1107. (2020).

Baker, J. C. et al. Oxygen isotopes in tree rings show good coherence between species and sites in Bolivia. Global Planet. Change 133, 298–308 (2015).

Barbour, M. M., & Farquhar, G. D. Relative humidity‐and ABA‐induced variation in carbon and oxygen isotope ratios of cotton leaves. Plant Cell Environ. 23, 473–485 (2000).

Barnard, R. L. et al. Evaporative enrichment and time lags between δ^18^O of leaf water and organic pools in a pine stand. Plant Cell Environ. 30, 539–550 (2007).

Baumgartrer, A., & Reichel, E. The world water balance; mean annual global, continental and maritime precipitacion, evaporation and run-off. (No. 628.1 B3). Elsevier Amsterdam (1975).

Begum, S., Nakaba, S., Oribe, Y., Kubo, T., & Funada, R. Changes in the localization and levels of starch and lipids in cambium and phloem during cambial reactivation by artificial heating of main stems of *Cryptomeria japonica* trees. Ann. Bot. 106, 885–895 (2010).

Berry, Z. C., Emery, N. C., Gotsch, S. G. & Goldsmith, G. R. Foliar water uptake: processes, pathways, and integration into plant water budgets. Plant Cell Environ. 42, 410–423 (2019).

Berry, Z. C., White, J. C. & Smith, W. K. Foliar uptake, carbon fluxes and water status are affected by the timing of daily fog in saplings from a threatened cloud forest. Tree Physiol. 34, 459–470 (2014).

Binks, O. et al. (2020). Canopy wetness in the Eastern Amazon, Agricultural and Forest Meteorology, 108250, https://doi.org/10.1016/j.agrformet.2020.108250

Boutton, T. W. Stable carbon isotope ratios of natural materials: I. Sample preparation and mass spectrometric analysis, *Carbon Isotope Techniques*, 1, 155–171, https://doi.org/10.1016/B978-0-12-179730-0.X5001-2 (Academic Press, 1991).

Brand, W. A. et al. Comprehensive inter-laboratory calibration of reference materials for δ^18^O versus VSMOW using various on-line high-temperature conversion techniques. Rapid Commun. Mass Sp. 23, 999–1019 (2009).

Breshears, D. D. et al. Foliar absorption of intercepted rainfall improves woody plant water status most during drought. Ecology 89, 41–47 (2008).

Burgess, S. S. O. & Dawson, T. E. The contribution of fog to the water relations of *Sequoia sempervirens* (D. Don): foliar uptake and prevention of dehydration. Plant Cell Environ. 27, 1023–1034 (2004).

Cassana, F. F., Eller, C. B., Oliveira, R. S. & Dillenburg, L. R. Effects of soil water availability on foliar water uptake of *Araucaria angustifolia*. Plant Soil 399, 147–157 (2016).

Cernusak, L. A. et al. Stable isotopes in leaf water of terrestrial plants. Plant Cell Environ. 39, 1087–1102 (2016).

Cernusak, L. A., Farquhar, G. D. & Pate, J. S. Environmental and physiological controls over oxygen and carbon isotope composition of Tasmanian blue gum, *Eucalyptus globulus*. Tree Physiol. 25, 129–146 (2005).

Choi, E. B. et al. Synchronization of tree-ring δ^18^O time series within and between tree species and provinces in Korea: A case study using dominant tree species in high elevations and archaeological wood. J. Wood Sci. 66, https://doi.org/10.1186/s10086-020-01901-3 (2020).

Craig, H. & Gordon L. I. Deuterium and oxygen-18 variations in the ocean and the marine atmosphere. In Proceedings of a Conference on Stable Isotopes in Oceanographic Studies and Palaeotemperatures (ed E. Tongiorgi), pp. 9–130. Lischi and Figli, Pisa (V. Lischi, 1965).

Cuntz, M., Ciais, P., Hoffmann, G. & Knorr, W. A comprehensive global three-dimensional model of δ^18^O in atmospheric CO_2_: 1. Validation of surface processes. J. Geophys. Res. 108, https://doi.org/10.1029/2002JD003153 (2003).

Dannoura, M. et al. The impact of prolonged drought on phloem anatomy and phloem transport in young beech trees. Tree Physiol. 39, 201–210 (2019).

Dansgaard, W. Stable isotopes in precipitation. Tellus 16, 436–468 (1964).

Dawson, T. E. Hydraulic lift and water use by plants: implications for water balance, performance and plant-plant interactions. Oecologia 95, 565–574 (1993).

Dawson, T. E., & Ehleringer, J. R. Streamside trees that do not use stream water. Nature 350, 335–337 (1991).

Dawson, T. E., & Goldsmith, G. R. The value of wet leaves. New Phytol. 219, 1156–1169 (2018).

De Wit, J. C., Van der Straaten, C. M. & Mook, W. G. Determination of the absolute hydrogen isotopic ratio of V-SMOW and SLAP. Geostandard. Newslett. 4, 33–36 (1980).

DeNiro, M. J. & Epstein, S. Relationship between the oxygen isotope ratios of terrestrial plant cellulose, carbon dioxide, and water. Science 204, 51–53 (1979).

Dole, M., Lane, G. A., Rudd, D. P. & Zaukelies, D. A. Isotopic composition of atmospheric oxygen and nitrogen. Geochim. Cosmochim. Ac. 6, 65–78 (1954).

Dongmann, G., Nurnberg, H. W., Förstel H. & Wagener K. On the enrichment of H_2_ ^18^O in the leaves of transpiring plants. Radiat. Environ. Bioph. 11, 41–52 (1974).

Dunin, F. X., O’loughlin, E. M. & Reyenga, W. Interception loss from eucalypt forest: lysimeter determination of hourly rates for long term evaluation. Hydrol. process. 2, 315–329 (1988).

Dunkerley, D. L. Evaporation of impact water droplets in interception processes: historical precedence of the hypothesis and a brief literature overview. J. Hydrol. 376, 599–604 (2009).

Eller, C. B., Lima, A. L. & Oliveira, R. S. Foliar uptake of fog water and transport belowground alleviates drought effects in the cloud forest tree species, *Drimys brasiliensis* (Winteraceae). New Phytol. 199, 151–162 (2013).

Ellsworth, P. V., & Sternberg, L. S. Biochemical effects of salinity on oxygen isotope fractionation during cellulose synthesis. New Phytol. 202, 784–789 (2014).

Epron, D., Dannoura, M., Ishida, A., & Kosugi, Y. Estimation of phloem carbon translocation belowground at stand level in a hinoki cypress stand. Tree Physiol. 39, 320–331 (2019).

Epstein, S., Yapp, C. J. & Hall, J. H. The determination of the D/H ratio of non-exchangeable hydrogen in cellulose extracted from aquatic and land plants. Earth Planet. Sc. Lett. 30, 241–251 (1976).

Farquhar, G. D., Barbour, M. M. & Henry, B. K. Interpretation of oxygen isotope composition of leaf material. In ‘Stable Isotopes: Integration of Biological, Ecological, and Geochemical Processes’. pp. 27–48. doi:10.1086/393961 Oxford (BIOS, 1998).

Farquhar, G. D. & Cernusak, L. A. On the isotopic composition of leaf water in the non-steady state. Funct. Plant Biol. 32, 293–303 (2005).

Farquhar G. D. & Gan, K. S. On the progressive enrichment of the oxygen isotopic composition of water along leaves. Plant Cell Environ. 26, 801–819 (2003).

Farquhar, G. D. & Lloyd, J. Carbon and oxygen isotope effects in the exchange of carbon dioxide between terrestrial plants and the atmosphere. In: Ehleringer J. R., Hall, A. E., and Farquhar, G. D., eds. Stable Isotopes and Plant Carbon-water Relations. 47–70, doi:10.1016/B978-0-08-091801-3.50011-8 San Diego (Academic Press, 1993).

Farquhar, G. D. et al. Vegetation effects on the isotope composition of oxygen in atmospheric CO_2_. Nature 363, 439–443 (1993).

Fernández, V. et al. Physico-chemical properties of plant cuticles and their functional and ecological significance. J. Exp. Bot. 68, 5293–5306 (2017).

Ferrio, J. P., Kurosawa, Y., Wang, M., & Mori, S. Hydraulic constraints to whole-tree water use and respiration in young Cryptomeria trees under competition. Forests 9, 449 (2018).

Gan, K. S., Wong, S. C. & Farquhar, G. D. Oxygen isotope analysis of plant water without extraction procedure, Handbook of Stable Isotope Analytical Techniques, 1, 473–481 https://doi.org/10.1016/B978-044451114-0/50023-5 (Elsevier, 2004).

Gessler, A. et al. Stable isotopes in tree rings: towards a mechanistic understanding of isotope fractionation and mixing processes from the leaves to the wood. Tree Physiol. 34, 796–818 (2014).

Gessler, A., Peuke, A. D., Keitel, C. & Farquhar, G. D. Oxygen isotope enrichment of organic matter in *Ricinus communis* during the diel course and as affected by assimilate transport. New Phytol. 174, 600–613 (2007).

Goldsmith, G. R., Matzke, N. J., & Dawson, T. E. The incidence and implications of clouds for cloud forest plant water relations. Ecol. Lett. 16, 307–314 (2013).

Goldsmith, G. R., Lehmann, M. M., Cernusak, L. A., Arend, M. & Siegwolf, R. T. Inferring foliar water uptake using stable isotopes of water. Oecologia 184, 763–766 (2017).

Gonfiantini, R., Gratsiu, S. & Tongiorgi, E. Oxygen isotopic composition of water in leaves. In Use of Isotopes and Radiation in Soil-Plant Nutrition Studies, pp.405–410 Vienna (IAEA, 1965).

Hanba, Y. T., Moriya, A. & Kimura, K. Effect of leaf surface wetness and wettability on photosynthesis in bean and pea. Plant Cell Environ. 27, 413–421 (2004).

Hannes et al. Diurnal variation in the isotope composition of plant xylem water biases the depth of root-water uptake estimates. Biogeosciences https://doi.org/10.5194/bg-2019-512 (2020).

Helle, G. & Panferov, O. Tree-Rings, isotopes, climate and environment: TRICE. Pages News 12, 22–23 (2004).

Helliker, B. R. & Ehleringer, J. R. Differential ^18^O enrichment of leaf cellulose in C3 versus C4 grasses, Funct. Plant Biol. 29, 435–442 (2002).

Helliker, B. R., & Richter, S. L. Subtropical to boreal convergence of tree-leaf temperatures, Nature 454, 511 (2008).

Hill, S. A., Waterhouse, J. S., Field, E. M., Switsur, V. R. & Ap Rees, T. Rapid recycling of triose phosphates in oak stem tissue. Plant Cell Environ. 18, 931–936 (1995).

Hörmann, G. et al. Calculation and simulation of wind controlled canopy interception of a beech forest in Northern Germany. Agr. Forest Meteorol. 79, 131–148 (1996).

Hoffmann, G. et al. A model of the Earth’s Dole effect. Global Biogeochem. Cy. 18, https://doi.org/10.1029/2003GB002059 (2004).

Horikawa, Y. & Sugiyama, J. Accessibility and size of Valonia cellulose microfibril studied by combined deuteration/rehydrogenation and FTIR technique, Cellulose 15, 419–424 (2008).

IAEA. Reference Sheet for International Measurement Standards, https://nucleus.iaea.org/rpst/documents/VSMOW_SLAP.pdf (2006).

Ingraham, N. L. Isotopic variations in precipitation, In Isotope tracers in catchment hydrology, 87–118, https://doi.org/10.1016/B978-0-444-81546-0.50010-0 (Elsevier, 1998).

Iida, S. I. et al. Intrastorm scale rainfall interception dynamics in a mature coniferous forest stand. J. Hydrol. 548, 770–783 (2017).

Kabeya, N., Katsuyama, M., Kawasaki, M., Ohte, N. & Sugimoto, A. Estimation of mean residence times of subsurface waters using seasonal variation in deuterium excess in a small headwater catchment in Japan. Hydrol. process. 21, 308–322 (2007).

Kagawa, A. & Leavitt, S. W. Stable carbon isotopes of tree rings as a tool to pinpoint the geographic origin of timber. J. Wood Sci. 56, 175–183 (2010).

Kagawa, A., Sano, M., Nakatsuka, T., Ikeda, T. & Kubo, S. An optimized method for stable isotope analysis of tree rings by extracting cellulose directly from cross-sectional laths. Chem. Geol. 393, 16–25 (2015).

Kagawa, A., Sugimoto, A. & Maximov, T. C. ^13^CO_2_ pulse-labelling of photoassimilates reveals carbon allocation within and between tree rings. Plant Cell Environ. 29, 1571–1584 (2006a).

Kagawa, A., Sugimoto, A. & Maximov, T. C. Seasonal course of translocation, storage and remobilization of ^13^C pulse-labeled photoassimilate in naturally growing *Larix gmelinii* saplings. New Phytol. 171, 793–804 (2006b).

Kagawa, A., Sugimoto, A., Yamashita, K. & Abe, H. Temporal photosynthetic carbon isotope signatures revealed in a tree ring through ^13^CO_2_ pulse-labelling. Plant Cell Environ. 28, 906–915 (2005).

Kim, K., & Lee, X. Transition of stable isotope ratios of leaf water under simulated dew formation. Plant, Cell Environ. 34, 1790–1801 (2011).

Klemm, O., Milford, C., Sutton, M. A., Spindler, G., & Van Putten, E. A climatology of leaf surface wetness. Theor. Appl. Climatol., 71, 107–117 (2002).

Koch, G. W., Sillett, S. C., Jennings, G. M., & Davis, S. D. The limits to tree height. Nature 428, 851–854 (2004).

Koch, K. Sucrose metabolism: regulatory mechanisms and pivotal roles in sugar sensing and plant development. Curr. Opin. Plant Biol. 7, 235–246 (2004).

Konter, O. et al. Climate sensitivity and parameter coherency in annually resolved δ^13^C and δ^18^O from *Pinus uncinata* tree-ring data in the Spanish Pyrenees. Chem. Geol. 377, 12–19 (2014).

Kottek, M., Grieser, J., Beck, C., Rudolf, B. & Rubel, F. World map of the Köppen-Geiger climate classification updated. Meteorol. Z. 15, 259–263 (2006).

Lacointe, A. Carbon allocation among tree organs: a review of basic processes and representation in functional-structural tree models. Ann. Forest Sci. 57, 521–533 (2000).

Laumer, W. et al. A novel approach for the homogenization of cellulose to use micro-amounts for stable isotope analyses. Rapid Commun. Mass Sp. 23, 1934–1940 (2009).

Laur, J. & Hacke, U. G. Exploring *Picea glauca* aquaporins in the context of needle water uptake and xylem refilling. New Phytol. 203, 388–400 (2014a).

Laur, J. & Hacke, U. G. The role of water channel proteins in facilitating recovery of leaf hydraulic conductance from water stress in *Populus trichocarpa*. PloS One 9, e111751 (2014b).

Leaney, F., Osmond, C., Allison, G. & Ziegler, H. Hydrogen-isotope composition of leaf water in C3 and C4 plants: its relationship to the hydrogen-isotope composition of dry matter. Planta 164, 215–220 (1985).

Lehmann, M. M. et al. The ^18^O-signal transfer from water vapour to leaf water and assimilates varies among plant species and growth forms. Plant Cell Environ. 43, 510–523 (2019).

Lehmann, M. M. et al. The effect of ^18^O-labelled water vapour on the oxygen isotope ratio of water and assimilates in plants at high humidity. New Phytol. 217, 105–116 (2018).

Lehmann, M. M. et al. More than climate: Hydrogen isotope ratios in tree rings as novel plant physiological indicator for stress conditions. Dendrochronologia 65, 125788 (2020).

Li, Z., Nakatsuka, T. & Sano, M. Tree-ring cellulose δ^18^O variability in pine and oak and its potential to reconstruct precipitation and relative humidity in central Japan. Geochem. J. 49, 125–137 (2015).

Limm, E. B., Simonin, K. A., Bothman, A. G. & Dawson, T. E. Foliar water uptake: a common water acquisition strategy for plants of the redwood forest. Oecologia 161, 449–459 (2009).

Loader, N. J. et al. Tree ring dating using oxygen isotopes: a master chronology for central England. J. Quat. Sci. 34, 475–490 (2019).

Luz, B., Barkan, E., Bender, M. L., Thiemens, M. H. & Boering, K. A. Triple-isotope composition of atmospheric oxygen as a tracer of biosphere productivity. Nature 400, 547–550 (1999).

McElrone, A. J., Choat, B., Gambetta, G. A. & Brodersen, C. R. Water uptake and transport in vascular plants. Nature Education Knowledge 4, 6 (2013).

Miyake, Y., Matsubaya, O., & Nishihara, C. An isotopic study on meteoric precipitation. Papers in Meteor. and Geophys. 19, 243–266 (1968).

Miyashita, A. et al. Long-term, short-interval measurements of the frequency distributions of the photosynthetically active photon flux density and net assimilation rate of leaves in a cool-temperate forest. Agr. Forest Meteorol. 152, 1–10 (2012).

Murakami, S. A proposal for a new forest canopy interception mechanism: Splash droplet evaporation. J. Hydrol. 319, 72–82 (2006).

Nabeshima, E., Nakatsuka, T., Kagawa, A., Hiura, T. & Funada, R. Seasonal changes of δD and δ^18^O in tree-ring cellulose of *Quercus crispula* suggest a change in post-photosynthetic processes during earlywood growth. Tree Physiol. 38, 1829–1840 (2018).

Nakada, R., Okada, N., Nakai, T., Kuroda, K., & Nagai, S. Water potential gradient between sapwood and heartwood as a driving force in water accumulation in wetwood in conifers. Wood Sci. Technol. 53, 407–424 (2019).

Nakai, W., Okada, N., Sano, M. & Nakatsuka, T. Sample preparation of ring-less tropical trees for δ^18^O measurement in isotope dendrochronology. Tropics 27, 49–58 (2018).

Nakatsuka, T. et al. Oxygen and carbon isotopic ratios of tree-ring cellulose in a conifer-hardwood mixed forest in northern Japan. Geochem. J. 38, 77–88 (2004).

Nakatsuka, T. et al. Reconstruction of multi-millennial summer climate variations in central Japan by integrating tree-ring cellulose oxygen and hydrogen isotope ratios. Clim. Past Discuss. https://doi.org/10.5194/cp-2020-6 (2020).

Nanko, K., Watanabe, A., Hotta, N. & Suzuki, M. Physical interpretation of the difference in drop size distributions of leaf drips among tree species. Agr. Forest Meteorol. 169, 74–84 (2013).

Newberry, S. L., Nelson, D. B., & Kahmen, A. Cryogenic vacuum artifacts do not affect plant water‐ uptake studies using stable isotope analysis. Ecohydrol. 10, e1892 (2017).

Offermann, C. et al. The long way down – are carbon and oxygen isotope signals in the tree ring uncoupled from canopy physiological processes? Tree Physiol. 31, 1088–1102 (2011).

Olsen, J., Seierstad, I., Vinther, B., Johnsen, S. & Heinemeier, J. Memory effect in deuterium analysis by continuous flow isotope ratio measurement. Int. J. Mass Spectrom. 254, 44–52 (2006).

Pfautsch, S., Renard, J., Tjoelker, M. G., & Salih, A. Phloem as capacitor: radial transfer of water into xylem of tree stems occurs via symplastic transport in ray parenchyma. Plant Physiol. 167, 963–971 (2015).

Qin, C., Yang, B., Braeuning, A., Grießinger, J., & Wernicke, J. Drought signals in tree-ring stable oxygen isotope series of Qilian juniper from the arid northeastern Tibetan Plateau. Global Planet. Change, 125, 48–59 (2015)

Ritpitakphong, U. et al. The microbiome of the leaf surface of *Arabidopsis* protects against a fungal pathogen. New Phytol. 210, 1033–1043 (2016).

Roden, J. Cross-dating of tree ring δ^18^O and δ^13^C time series. Chem. Geol. 252, 72–79 (2008).

Roden, J. S., & Ehleringer, J. R. Hydrogen and oxygen isotope ratios of tree-ring cellulose for riparian trees grown long-term under hydroponically controlled environments. Oecologia 121, 467–477 (1999a).

Roden, J. S. & Ehleringer, J. R. Observations of hydrogen and oxygen isotopes in leaf water confirm the Craig-Gordon model under wide-ranging environmental conditions. Plant Physiol. 120, 1165–1174 (1999b).

Roden, J. S., Johnstone, J. A. & Dawson, T. E. Intra-annual variation in the stable oxygen and carbon isotope ratios of cellulose in tree rings of coast redwood (*Sequoia sempervirens*). The Holocene 19, 189–197 (2009).

Roden, J. S., Johnstone, J. A. & Dawson, T. E. Regional and watershed-scale coherence in the stable-oxygen and carbon isotope ratio time series in tree rings of coast redwood (*Sequoia sempervirens*). Tree-Ring Res. 67, 71–86 (2011).

Roden, J. S., Lin, G. & Ehleringer, J. R. A mechanistic model for interpretation of hydrogen and oxygen isotope ratios in tree-ring cellulose. Geochim. Cosmochim. Ac. 64, 21–35 (2000).

Rossi, S. et al. Conifers in cold environments synchronize maximum growth rate of tree-ring formation with day length. New Phytol. 170, 301–310 (2006).

Schmidt, H. L., Werner, R. A. & Roßmann, A. ^18^O pattern and biosynthesis of natural plant products. Phytochemistry 58, 9–32 (2001).

Schoeller, D. A., Van Santen, E. Measurement of energy expenditure in humans by doubly labeled water method. J. Appl. Physiol. 53, 955–959 (1982).

Schreel, J. D. et al. Identifying the pathways for foliar water uptake in beech (*Fagus sylvatica* L.): a major role for trichomes. Plant J. 103, 769–780 (2020).

Schreel, J. D. M. & Steppe, K. Foliar water uptake in trees: negligible or necessary? Trends Plant Sci. https://doi.org/10.1016/j.tplants.2020.01.003 (2020).

Seiler, J. R. & Johnson, J. D. Physiological and morphological responses of three half-sib families of loblolly pine to water-stress conditioning. Forest Sci. 34, 487–495 (1988).

Seo, J. W. et al. Oxygen isotope ratios of subalpine conifers in Jirisan National Park, Korea and their dendroclimatic potential. Dendrochronologia 57, 125626 (2019).

Sevanto, S., Hölttä, T., & Holbrook, N. M. Effects of the hydraulic coupling between xylem and phloem on diurnal phloem diameter variation. Plant Cell Environ. 34, 690–703 (2011).

Shestakova, T. A., Aguilera, M., Ferrio, J. P., Gutiérrez, E., & Voltas, J. Unravelling spatiotemporal tree-ring signals in Mediterranean oaks: a variance–covariance modelling approach of carbon and oxygen isotope ratios. Tree Physiol., 34, 819–838 (2014).

Singer, M. B. et al. Contrasting water-uptake and growth responses to drought in co-occurring riparian tree species. Ecohydrology 6, 402–412 (2013).

Smith, J. A. C. & Milburn, J. A. Phloem turgor and the regulation of sucrose loading in *Ricinus communis* L. Planta 148, 42–48 (1980).

Song, X., Farquhar, G. D., Gessler, A., & Barbour, M. M. Turnover time of the non◻structural carbohydrate pool influences δ^18^ O of leaf cellulose. Plant, Cell Environ. 37, 2500–2507 (2014).

Speakman, J. Doubly labelled water: theory and practice. Springer Science & Business Media (1997).

Steppe, K. et al. Direct uptake of canopy rainwater causes turgor-driven growth spurts in the mangrove *Avicennia marina*. Tree Physiol. 38, 979–991 (2018).

Sternberg, L. D. S. L. O. R. Oxygen stable isotope ratios of tree-ring cellulose: The next phase of understanding. New Phytol. 181, 553–562 (2009).

Sternberg, L. D. S. L., Anderson, W. T. & Morrison, K. Separating soil and leaf water ^18^O isotopic signals in plant stem cellulose. Geochim. Cosmochim. Ac. 67, 2561–2566 (2003).

Sternberg, L. & DeNiro, M. J. Isotopic composition of cellulose from C3, C4, and CAM plants growing near one another. Science 220, 947–949 (1983).

Sternberg, L. D. S., Deniro, M. J. & Savidge, R. A. Oxygen isotope exchange between metabolites and water during biochemical reactions leading to cellulose synthesis. Plant Physiol. 82, 423–427 (1986).

Sternberg, L., Pinzon, M. C., Anderson, W. T. & Jahren, A. H. Variation in oxygen isotope fractionation during cellulose synthesis: intramolecular and biosynthetic effects. Plant Cell Environ. 29, 1881–1889 (2006).

Terwilliger, V. J. & Deniro, M. J. Hydrogen isotope fractionation in wood-producing avocado seedlings: Biological constraints to paleoclimatic interpretations of δD values in tree ring cellulose nitrate. Geochim. Cosmochim. Ac. 59, 5199–5207 (1995).

Treydte, K. et al. Seasonal transfer of oxygen isotopes from precipitation and soil to the tree ring: source water versus needle water enrichment. New Phytol. 202, 772–783 (2014).

Tsujimura, M. & Tanaka, T. Evaluation of evaporation rate from forested soil surface using stable isotopic composition of soil water in a headwater basin. Hydrol. Process. 12, 2093–2103 (1998).

Vesala, T. et al. Effect of leaf water potential on internal humidity and CO_2_ dissolution: reverse transpiration and improved water use efficiency under negative pressure. Front. Plant Sci. 8, 54 (2017).

Wang, X., Xiao, H., Cheng, Y. & Ren, J. Leaf epidermal water-absorbing scales and their absorption of unsaturated atmospheric water in *Reaumuria soongorica*, a desert plant from the northwest arid region of China. J. Arid Environ. 128, 17–29 (2016).

Waterhouse, J. S., Switsur, V. R., Barker, A. C., Carter, A. H. C. & Robertson, I. Oxygen and hydrogen isotope ratios in tree rings: how well do models predict observed values? Earth Planet. Sc. Lett. 201, 421–430 (2002).

Welp, L. R. et al. Interannual variability in the oxygen isotopes of atmospheric CO_2_ driven by El Nino. Nature 477, 579–582 (2011).

Wershaw, R. L., Friedman, I., Heller, S. J. & Frank, P. A. Hydrogen isotopic fractionation of water passing through trees. In Advances in Organic Geochemistry (ed. G.D. Hobson), International Series of Monographs in Earth Sciences 32, pp. 55–67 (Pergamon Press, 1966)

Windt, C. W., Vergeldt, F. J., De Jager, P. A., & Van As, H. MRI of long◻distance water transport: a comparison of the phloem and xylem flow characteristics and dynamics in poplar, castor bean, tomato and tobacco. Plant Cell Environ. 29, 1715–1729 (2006).

Yakir, D., Berry, J. A., Giles, L. & Osmond, C. B. Isotopic heterogeneity of water in transpiring leaves: identification of the component that controls the δ^18^O of atmospheric O_2_ and CO_2_. Plant Cell Environ. 17, 73–80 (1994).

Yakir, D., & DeNiro, M. J. Oxygen and hydrogen isotope fractionation during cellulose metabolism in *Lemna gibba* L. Plant Physiol. 93, 325–332 (1990).

Yakir, D., DeNiro, M. J. & Gat, J. R. Natural deuterium and oxygen-18 enrichment in leaf water of cotton plants grown under wet and dry conditions: evidence for water compartmentation and its dynamics. Plant Cell Environ. 13, 49–56 (1990).

Yakir, D., DeNiro, M. J. & Rundel, P. W. Isotopic inhomogeneity of leaf water: evidence and implications for the use of isotopic signals transduced by plants. Geochim. Cosmochim. Ac. 53, 2769–2773 (1989).

Yan, X. et al. Molecular mechanisms of foliar water uptake in a desert tree. AoB Plants 7, https://doi.org/10.1093/aobpla/plv129 (2015).

Yoshida, R. et al. An application of a physical vegetation model to estimate climate change impacts on rice leaf wetness, J. Appl. Meteorol. Clim. 54, 1482–1495 (2015).

Zhao, L. et al. Significant difference in hydrogen isotope composition between xylem and tissue water in *Populus euphratica*. Plant Cell Environ. 39, 1848–1857 (2016).

